# Detecting genome-wide directional effects of transcription factor binding on polygenic disease risk

**DOI:** 10.1101/204685

**Authors:** Yakir A Reshef, Hilary K Finucane, David R Kelley, Alexander Gusev, Dylan Kotliar, Jacob C Ulirsch, Farhad Hormozdiari, Joseph Nasser, Luke O’Connor, Bryce van de Geijn, Po-Ru Loh, Shari Grossman, Gaurav Bhatia, Steven Gazal, Pier Francesco Palamara, Luca Pinello, Nick Patterson, Ryan P Adams, Alkes L Price

## Abstract

Biological interpretation of GWAS data frequently involves analyzing unsigned genomic annotations comprising SNPs involved in a biological process and assessing enrichment for disease signal. However, it is often possible to generate signed annotations quantifying whether each SNP allele promotes or hinders a biological process, e.g., binding of a transcription factor (TF). Directional effects of such annotations on disease risk enable stronger statements about causal mechanisms of disease than enrichments of corresponding unsigned annotations. Here we introduce a new method, signed LD profile regression, for detecting such directional effects using GWAS summary statistics, and we apply the method using 382 signed annotations reflecting predicted TF binding. We show via theory and simulations that our method is well-powered and is well-calibrated even when TF binding sites co-localize with other enriched regulatory elements, which can confound unsigned enrichment methods. We further validate our method by showing that it recovers known transcriptional regulators when applied to molecular QTL in blood. We then apply our method to eQTL in 48 GTEx tissues, identifying 651 distinct TF-tissue expression associations at per-tissue FDR < 5%, including 30 associations with robust evidence of tissue specificity. Finally, we apply our method to 46 diseases and complex traits (average *N* = 289,617) and identify 77 annotation-trait associations at per-trait FDR < 5% representing 12 independent TF-trait associations, and we conduct gene-set enrichment analyses to characterize the underlying transcriptional programs. Our results implicate new causal disease genes (including causal genes at known GWAS loci), and in some cases suggest a detailed mechanism for a causal gene’s effect on disease. Our method provides a new way to leverage functional data to draw inferences about disease etiology.

## Introduction

Mechanistic interpretation of GWAS data sets has become a central challenge for efforts to learn about the biological underpinnings of disease. One successful paradigm for such efforts has been GWAS enrichment, in which a genome annotation containing SNPs that affect some biological process is shown to be enriched for GWAS signal^1–7^. However, there are instances in which experimental data allow us not only to identify SNPs that affect a biological process, but also to predict which SNP alleles promote the process and which SNP alleles hinder it, thereby enabling us to assess whether there is a systematic association between SNP alleles’ direction of effect on the process and their direction of effect on a trait. Transcription factor (TF) binding, which plays a major role in human disease^1,8–12^, represents an important case in which such signed functional annotations are available: because TFs have a tendency to bind to specific DNA sequences, it is possible to estimate whether the sequence change introduced by a SNP allele will increase or decrease binding of a TF^1,13–19^.

Detecting genome-wide directional effects of TF binding on disease would constitute a significant advance in terms of both evidence for causality and understanding of biological mechanism. Regarding causality, this is because directional effects are not confounded by simple co-localization in the genome (e.g., of TF binding sites with other regulatory elements), and thus provide stronger evidence for causality than is available using unsigned enrichment methods. Regarding biological mechanism, it is currently unknown whether disease-associated TFs affect only a few disease genes or whether transcriptional programs comprising many target genes are responsible for TF associations; a genome-wide directional effect implies the latter model (see Discussion).

Here we introduce a new method, signed LD profile (SLDP) regression, for quantifying the genome-wide directional effect of a signed functional annotation on polygenic disease risk, and apply it in conjunction with 382 annotations each reflecting predicted binding of a particular TF in a particular cell line. Our method requires only GWAS summary statistics^20^, accounts for linkage disequilibrium and untyped causal SNPs, and is computationally efficient. We validate the method via extensive simulations, including null simulations confounded by unsigned enrichment as might arise from the co-localization of TF binding sites with other regulatory elements^5,13^. We further validate the method by applying it to molecular QTL in blood^21^ and showing that it recovers known transcriptional regulators. We then apply the method to eQTL in 48 tissues from the GTEx consortium^22^ and to 46 diseases and complex traits, demonstrating genome-wide directional effects of TF binding in both settings. We further characterize the transcriptional programs underlying our complex trait associations via gene-set enrichment analyses using gene sets from the Molecular Signatures Database^23,24^ (MSigDB).

## Results

### Overview of methods

Our method for quantifying directional effects of signed functional annotations on disease risk, signed LD profile regression, relies on the fact that the signed marginal association of a SNP to disease includes signed contributions from all SNPs tagged by that SNP. Given a signed functional annotation with a directional linear effect on disease risk, the vector of marginal SNP effects on disease risk will therefore be proportional (in expectation) to a vector quantifying each SNP’s aggregate tagging of the signed annotation, which we call the *signed LD profile* of the annotation. Thus, our method detects directional effects by assessing whether the vector of marginal SNP effects and the signed LD profile are systematically correlated genome-wide.

More precisely, under a polygenic model^25^ in which true causal SNP effects are correlated with a signed functional annotation, we show that

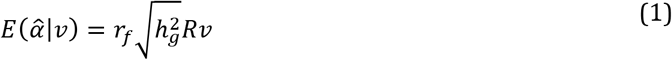

where 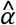 is the vector of marginal correlations between SNP alleles and a trait, *ν* is the signed functional annotation (re-scaled to norm 1) reflecting, e.g., the signed effect of a SNP on TF binding, *R* is the LD matrix, 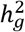 is the SNP-heritability of the trait, and *r_f_* is the correlation between the vector *ν* and the vector of true causal effects of each SNP, which we call the *functional correlation.* (*r_f_* can be interpreted as a form of genetic correlation; the value of 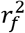 cannot exceed the proportion of SNP-heritability explained by SNPs with non-zero values of *ν*.) Equation (1), together with an estimate of 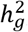, allows us to estimate *r_f_* by regressing 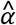 on the signed LD profile *Rν* of *ν*. We assess statistical significance by randomly flipping the signs of entries of *ν*, with consecutive SNPs being flipped together in large blocks (e.g., ~300 blocks total), to obtain a null distribution and corresponding P-values and false discovery rates (FDRs). To improve power, we use generalized least-squares regression, incorporating weights to account for the fact that SNPs in linkage disequilibrium (LD) provide redundant information due to their correlated values of 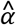. We remove the major histocompatibility complex (MHC) region from all analyses due to its unusual LD patterns. We perform a multiple regression that explicitly conditions on a “signed background model” corresponding to directional effects of minor alleles in five equally sized minor allele frequency (MAF) bins, which could reflect confounding due to genome-wide negative selection or population stratification. We note that signed LD profile regression requires signed effect size estimates 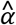 and quantifies directional effects, in contrast to stratified LD score regression^5^, which analyzes unsigned *χ*^2^ statistics and quantifies unsigned heritability enrichment. Details of the method are described in the Online Methods section and the Supplementary Note; we have released open-source software implementing the method (see URLs).

We applied signed LD profile regression using a set of 382 signed annotations *ν*, each quantifying the predicted effects of SNP alleles on binding of a particular TF in a particular cell line. We constructed the annotations by training a sequence-based neural network predictor of ChIP-seq peak calls, using the Basset software^19^, to predict the results of 382 TF binding ChIP-seq experiments from ENCODE^26^ and comparing the neural network’s predictions for the major and minor allele of each SNP in the ChIP-seq peaks. The 382 experiments spanned 75 distinct TFs and 84 distinct cell lines. Because each annotation contained non-zero entries only for SNPs lying inside ChIP-seq peaks of the corresponding ChIP-seq experiment, the resulting annotations were sparse, with only 0.2% of SNPs having nonzero entries on average (see Online Methods and Table S1).

### Simulations

We performed simulations with real genotypes, simulated phenotypes, and our 382 signed TF binding annotations to assess null calibration, robustness to confounding, and power. All simulations used well-imputed genome-wide genotypes from the GERA cohort^27^, corresponding to *M* = 2.7 million SNPs and *N* = 47,360 individuals of European ancestry. We simulated traits using normally distributed causal effect sizes (with annotation-dependent mean and variance in some cases), with 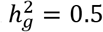. Further details of the simulations are provided in the Online Methods section.

We first performed null simulations involving a heritable trait with no unsigned enrichment or directional association to any of our 382 annotations. In 1,000 independent simulations, we applied signed LD profile regression to test each of our 382 annotations for a directional effect. The resulting P-values were well-calibrated (see Figure 1a and Table S2). Analyses of the P-value distribution for each annotation in turn confirmed correct calibration for these annotations (see Figure S1a).

**Figure 1.**
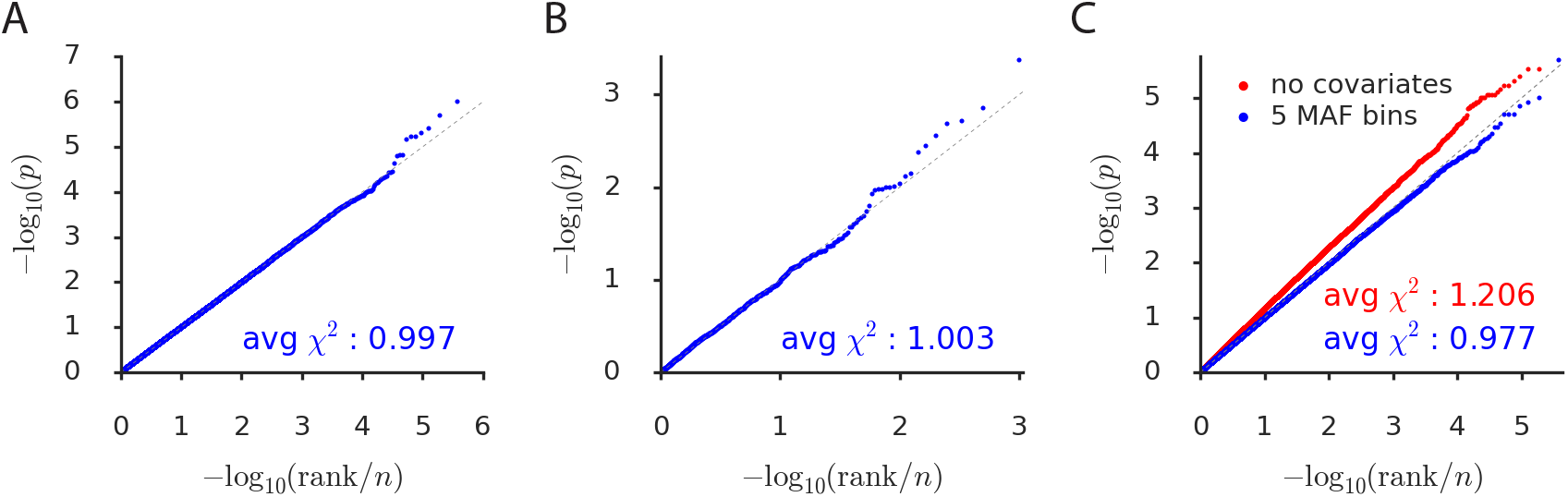
Simulations assessing null calibration. We report null calibration (q-q plots of –log_10_(*p*) values) in simulations of (a) no enrichment, (b) unsigned enrichment, and (c) directional effects of minor alleles. The q-q plots are based on (a) 382 annotations × 1,000 simulations = 382,000, (b) 1,000, and (c) two sets of 382 × 1,000 = 382,000 P-values. A 5-MAF-bin signed background model is included in all cases except for the red points in part (c), which are computed with no covariates. We also report the average *χ*^2^-statistic corresponding to each set of P-values. Numerical results are reported in Table S2.

We next performed null simulations involving a trait with unsigned enrichment but no directional effects; these simulations were designed to mimic unsigned genomic confounding in which the binding sites of some TF lie in or near regulatory regions that are enriched for heritability for reasons other than binding of that TF. In 1,000 independent simulations, we randomly selected an annotation, simulated a trait in which the annotation had a 20x unsigned enrichment^5^ (but no directional effect), and applied signed LD profile regression to test the annotation for a directional effect. We again observed well-calibrated P-values (see Figure 1b). It is notable that our method is well-calibrated even though it has no knowledge of the unsigned genomic confounder; this contrasts with unsigned enrichment approaches such as heritability partitioning, in which unsigned genomic confounders must be carefully accounted for and modeled^5^.

We next performed null simulations to assess whether our method remains well-calibrated in the presence of confounding due to genome-wide directional effects of minor alleles on both disease risk and TF binding, which could arise due to genome-wide negative selection or population stratification. We simulated a trait for which 10% of heritability is explained by directional effects of minor alleles in the bottom fifth of the MAF spectrum (roughly MAF < 5%). In 1,000 independent simulations, we applied signed LD profile regression to test each of our 382 annotations for a directional effect. P-values were well-calibrated for the default version of the method, which conditions on the 5-MAF-bin signed background model, but were not well-calibrated without conditioning on this model (see Figure 1c). (We note that this represents a best-case scenario in which the background model exactly matches the confounding being simulated, up to differences in MAF between the reference panel and the GWAS sample, and we caution that our method may not be appropriate for annotations with much stronger correlations to minor alleles than the annotations that we analyze here; see Figure S1b.) The incorrect calibration that we observe when we do not include our signed background model could potentially be explained by genome-wide negative selection against decreased TF binding^28^, which would result in a bias in the sign of the entries of our annotations. Indeed, most of our annotations show a small but highly significant bias of minor alleles toward decreasing TF binding (see Figure S2) that is consistent with this explanation; however, it is also possible that this bias is a result of our procedure for constructing the annotations, and we do not explore it further in this work. To ameliorate potential confounding by directional effects of minor alleles, we condition on the signed background model in all analyses in this paper unless stated otherwise.

Finally, we performed causal simulations with true directional effects to assess the power and establish the unbiasedness of signed LD profile regression. At default parameter settings, the method is well-powered to detect directional effects corresponding to a functional correlation of 2-6% (see Figure 2a and Table S3), similar to values observed in analyses of real traits (see below). Notably, the power of the method is improved dramatically by our use of generalized least-squares to account for redundant information (see Figure 2a). Our method is also much more powerful than a naive method that regresses the vector of GWAS summary statistics on the annotation rather than its signed LD profile, an approach that does not model untyped causal SNPs in linkage disequilibrium with typed SNPs (see Figure S3). The power of our method increases with sample size and SNP-heritability (see Figure S4), and is only minimally affected by within-Europe reference panel mismatch (see Figure S5). In all instances, our method produced either unbiased or nearly unbiased estimates of functional correlation and related quantities (see Figure 2b and Figure S6).

**Figure 2.**
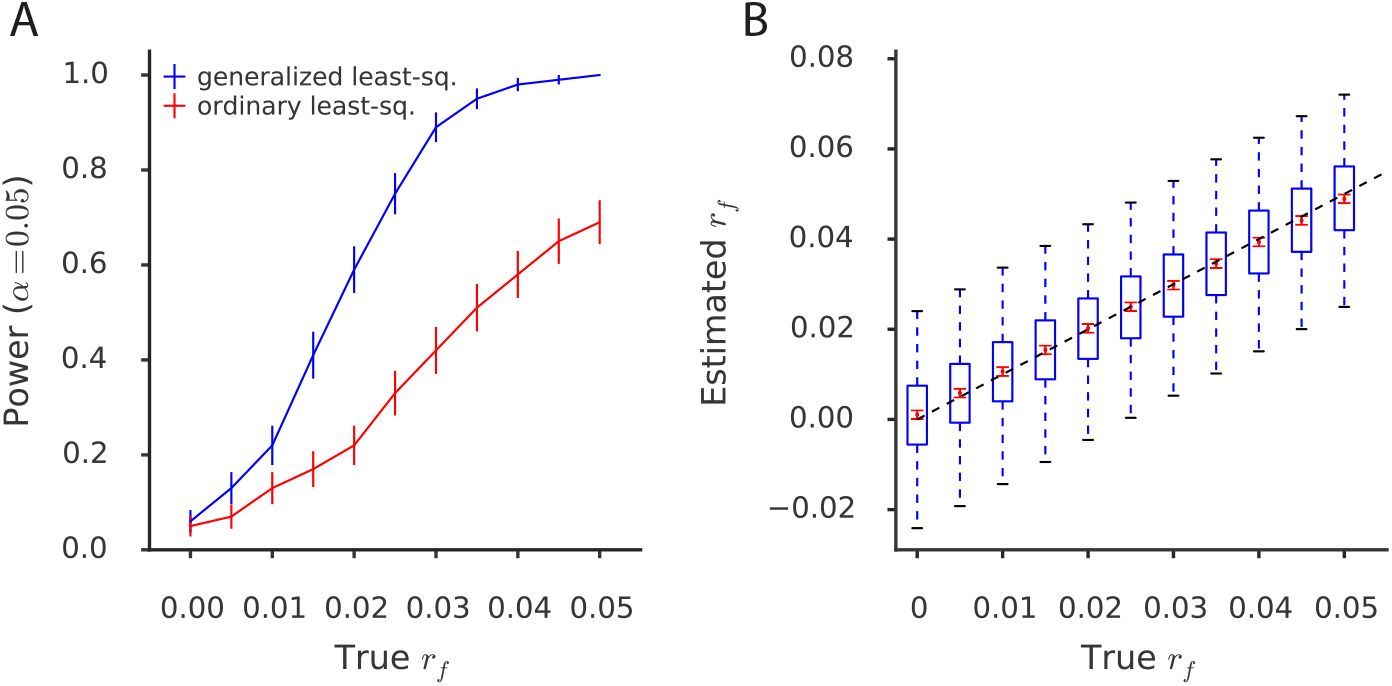
Simulations assessing power, bias, and variance. (a) Power curves under simulation scenarios comparing signed LD profile regression using generalized least-squares (i.e., weighting) to an ordinary (i.e., unweighted) regression of the summary statistics on the signed LD profile. Error bars indicate standard errors of power estimates. (b) Assessment of bias and variance of the signed LD profile regression estimate of *r_f_* at realistic sample size (47,360) and heritability (0.5), across a range of values of the true *r_f_*. Blue box and whisker plots depict the sampling distribution of the statistic, while the red dots indicate the estimated sample mean and the red error bars indicate the standard error around this estimate. Numerical results are reported in Table S3.

### Analysis of molecular traits in blood

TF binding is known to affect gene expression and other molecular traits^29^, and regulatory relationships in blood are particularly well-characterized^30^. We therefore applied signed LD profile regression to 12 molecular traits in blood with an average sample size of *N* = 149, to further validate the method. We first analyzed cis-eQTL data based on RNA-seq experiments in three blood cell types from the BLUEPRINT consortium^21^ (see Online Methods). For each cell type, we collapsed eQTL summary statistics across 15,023-17,081 genes into a single vector of summary statistics for aggregate expression by meta-analyzing, for each SNP, the marginal effect sizes of that SNP for the expression of all nearby genes (within 500kb; see Online Methods and Table S4).

We tested each of our 382 TF binding annotations for a directional effect on aggregate expression in each of the three blood cell types. We detected a total of 409 significant associations at a per-trait FDR of 5% (36% of annotation-blood cell type expression pairs tested) representing 107 distinct TF-blood cell type expression associations (see Figure 3a and Table S5a; P-values from ≤ 10^−6^ to 2.0×10^−2^). All of the detected associations were positive, implying that greater binding of these TFs leads to greater expression (in aggregate across genes) and matching the known tendency of TF binding to promote rather than repress transcription for many TFs^29^. Indeed, 170 of our 382 annotations (45%) correspond to TFs annotated as having activating activity and no repressing activity in UniProt^31^ (“activating”) and 174 (46%) correspond to TFs annotated as having either both activating and repressing activity (“ambiguous”); in contrast, only 38 (10%) correspond to TFs annotated as having repressing activity and no activating activity (“repressing”).

**Figure 3.**
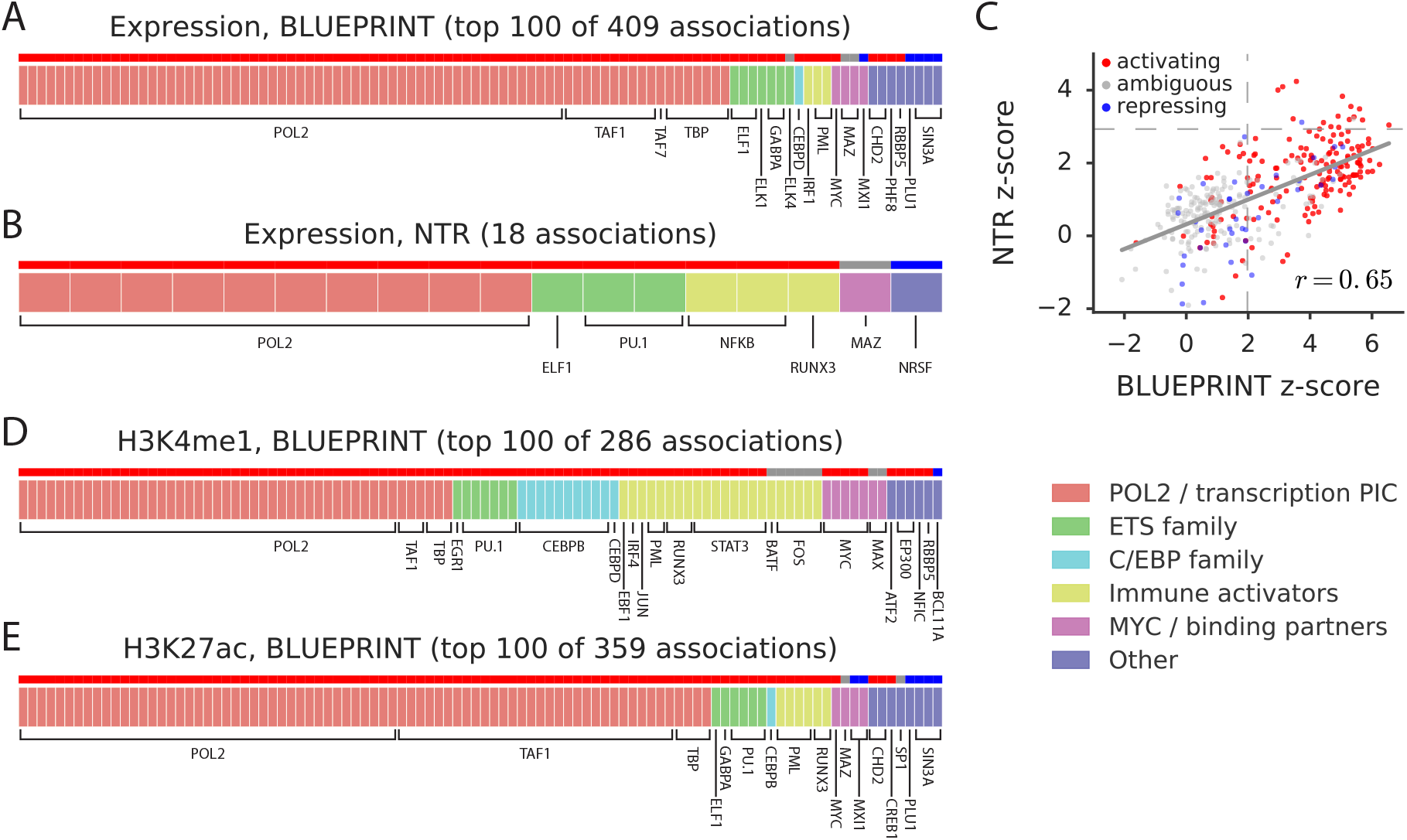
Analysis of blood molecular traits using signed LD profile regression. Each segmented bar in (a,b,d,e) represents the set of significant annotations (or top 100 associations) at a per-trait FDR of 5% for the indicated traits, with each annotation corresponding to a particular TF profiled in a particular cell line. Results in (a,d,e) are aggregated across the 3 BLUEPRINT cell types. The stripe above each segmented bar is colored red for UniProt (unambiguously) “activating” TFs (170 of 382 annotations; see main text), gray for “ambiguous” TFs (174 of 382 annotations), and blue for (unambiguously) “repressing” TFs (38 of 382 annotations). We note that the large number of associations detected in this analysis is expected due to widespread effects of TF binding on gene expression and chromatin as well as the strong representation of transcriptional activators among our annotations (see Table S1). (c) z-scores from the analyses of expression in the NTR data set and neutrophil expression in the BLUEPRINT data set, respectively, for each of the 382 annotations tested; red, gray, and blue again indicate UniProt (unambiguously) “activating” TFs, “ambiguous” TFs, and (unambiguously) “repressing” TFs, respectively. Dashed lines represent significance thresholds for 5% FDR. Numerical results are reported in Table S5.

As expected, many of the associations that we detected recapitulate known aspects of transcriptional regulation. For example, the most strongly associated TF binding annotations included RNA polymerase II in many cell lines, along with the two other profiled members of the transcription pre-initiation complex (PIC), TATA-associated Factor 1 (TAF1) and TATA Binding Protein (TBP). We also detected associations for TFs unrelated to the PIC but known to have activating activity, such as the ETS family members GABPA, ELF1, and ELK1^32^, as well as the immune- and cancer-related transcriptional activators interferon regulatory factor 1 (IRF1) and promyelocytic leukemia protein (PML)^33,34^. Overall, the majority of the positive associations (318 out of 409; 78%) involved (unambiguously) activating TFs (compared with 170 of our 382 (45%) annotations; *P* = 7.0×10^−43^ for difference using one-sided binomial test; see Figure 3a and Online Methods). 196 of the 409 associations replicated (same direction of effect with nominal *P* < 0.05) in an independent set of whole-blood eQTL summary statistics based on expression array experiments from the Netherlands Twin Registry (NTR)^35^, including all of the examples mentioned above except IRF1 (see Figure 3b and Table S5b). Across all 382 annotations analyzed, we observed a correlation of *r* = 0.65 between z-scores for signed annotation effects in the BLUEPRINT neutrophil and NTR data sets (see Figure 3c and Table S5c).

We next conducted a similar analysis using histone QTL (H3K27me1 and H3K27ac) and methylation QTL from the BLUEPRINT data set. We detected 645 significant associations at a per-trait FDR of 5% (28% of annotation-blood cell type QTL pairs tested), four of which were negative. These results included 286 significant associations for H3K27me1 QTL, 359 for H3K27ac QTL, and 0 for methylation QTL (79, 98, and 0 distinct TF-cell type QTL associations, respectively; see Figure 3d,e and Table S5d,e; P-values from ≤ 10^−6^ to 1.9×10^−2^). Once again, many of the detected associations recover known aspects of histone mark biology, as expected. For example, the TFs most strongly associated to H3K4me1 included PU.1 and CEBPB, both of which act to increase H3K4me1 in blood cells and play strong roles in differentiation of those cell types^36–39^, and binding of MYC, which has a known role as a chromatin modifier^40,41^, including of H3K4 methylation^42^. We also observed a strong positive association between H3K27ac and CREB1, a binding partner of the lysine acetyltransferase EP300, as well as a weaker positive association for EP300 itself, matching the well-documented role of both factors in creation and maintenance of this mark^43,44^. Several of our positive associations, such as the associations detected for the ETS TFs (including PU.1), are also consistent with a prior study^45^ that detected correlations between changes in position-weight matrix scores induced by SNPs and allelic imbalance at those SNPs in ChIP-seq data for these marks. The four negative associations that we detected involved MAFK and MAFF, both of which lack a transactivation domain^46^, as well as CTCF, which is known to act as an insulator^47,48^. Once again, the majority of the positive associations (528 out of 641; 82%) involved (unambiguously) activating TFs^31^ (onesided binomial *P* = 1.9×10^−9^).

### Analysis of gene expression across 48 GTEx tissues

We next applied signed LD profile regression to GTEx eQTL across 48 tissues^22^ (average *N* = 214) in order to draw inferences about transcriptional regulation across these tissues, including tissue-specific regulatory effects. We first tested each of our 382 TF binding annotations for a directional effect on expression in each of the 48 tissues in turn, analogous to our previous analysis of molecular traits in blood. For each significant association that we detected, we then assessed the association for tissue specificity by checking whether it remained at least as significant when conditioning on average eQTL effects across tissues (see Online Methods and Table S6). This criterion for tissue-specificity is conservative and stands in contrast to, e.g., reporting associations that remain significant at a specified threshold after conditioning. The latter approach is susceptible to the fact that conditioning on a noisily measured confounder can produce false positives^49^; associations meeting the former criterion are likely to be robustly tissue-specific.

Our analysis yielded 2,330 annotation-tissue expression associations at a per-trait FDR of 5% (13% of annotation-tissue expression pairs tested), representing 651 distinct TF-tissue expression associations of which 30 were robustly tissue-specific in our conditional analysis (see Figure 4 and Table S7). We detected both known and novel associations. The known TF-tissue associations that we detected include: activating roles for FOXA1 and FOXA2 in pancreas and other gastrointestinal tissues, recapitulating the well-known master regulatory function played by these factors in these tissues^50–52^; an activating role for early B-cell factor 1 (EBF-1) in lymphocytes^53,54^; an activating role for hepatocyte nuclear factor 4*γ* (HNF4G) – and a tissue-specific activating role for the related protein HNF4A – in liver^55,56^; a tissue-specific activating role for PU.1 in spleen, an organ that is hyperplastic when the *PU.1* gene is virally perturbed in mice^57^; and tissue-specific activating roles for FOS in fibroblasts, the animal tissue in which FOS was originally discovered^58^, as well as in nerve tissue, a tissue in which FOS deficiency causes numerous abnormalities^59–61^. Our results for these transcription factors contrast with the ubiquitous activating signatures detected for the three profiled transcription factors comprising the transcription pre-initiation complex (PIC; see above), POL2, TAF1, and TBP, for which we detected significant positive associations in 33 of the 48 tissues (69%) and 89% of the 28 tissues with a sample size above 150. Our results were concordant with absolute gene expression measurements of the detected TFs in the associated GTEx tissue samples: the proportion of significant TF associations in which the TF was expressed above a minimum threshold in the associated GTEx tissue (see Online Methods) was greater than the corresponding proportion for non-significant TFs in 32 out of the 34 tissues for which we could perform the comparison (*p* = 2.1×10^−15^ for trend across tissues; see Figure S7 for breakdown by tissue).

**Figure 4.**
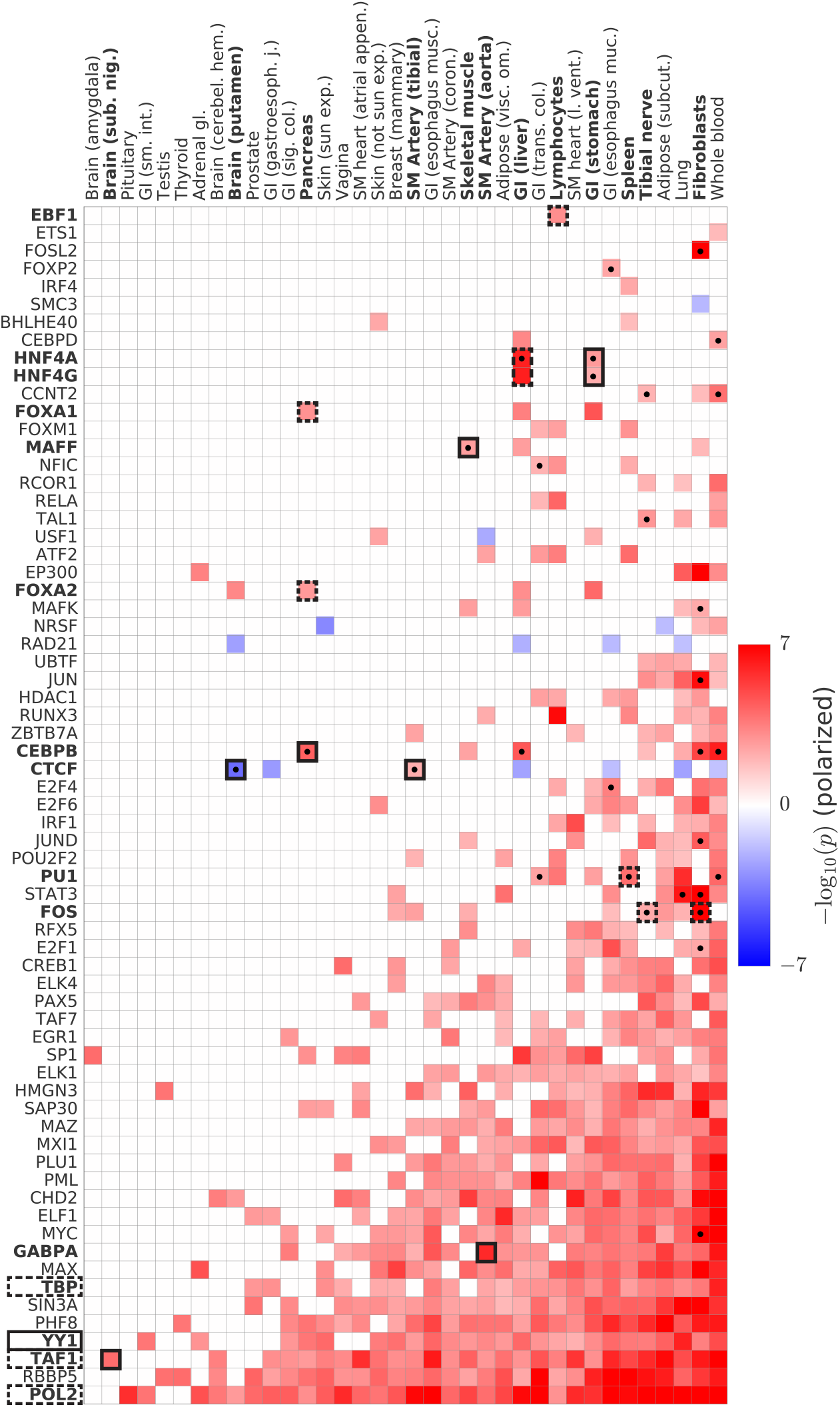
Analysis of GTEx eQTLs using signed LD profile regression. We plot polarized -log_10_ (*p*) values for all significant associations as a heatmap. Columns denote the 36 GTEx tissues (of 48 GTEx tissues tested) with significant associations. Rows denote the 67 TFs (of 75 TFs tested) with significant associations, collapsing all annotations corresponding to a single TF into one row and displaying in each case the most significant result. Cells with dots indicate associations that show robust evidence for tissue-specificity in our conditional analysis (see main text). Cells indicated in outline correspond to associations described in the main text, with dashed outline indicating known associations and solid outline indicating previously unknown associations or associations supporting emerging theories. Numerical results are reported in Table S7.

Our analysis also uncovered many previously unknown associations in less well-studied tissues that support emerging theories of disease. For example, the most significant association that we detected in aorta is a previously unreported activating role for GABPA. Though the regulatory role of this transcription factor in aorta has not been experimentally studied in detail, it is one of several related TFs whose binding sites were reported to be enriched near genes that were differentially expressed in aortic aneurysm samples vs. control samples^62^. Our association therefore provides direct evidence for the relevance of this TF to *in vivo* aortic gene regulation, as well as potential insight into the underlying mechanism behind aortic aneurysm. In addition, our top — and only — association in the brain tissue substantia nigra is TAF1. Neurodegeneration in the substantia nigra is a hallmark of Parkinson’s disease^63^ and TAF1 was proven earlier this year (through detailed experimental work) to be the causal gene in a rare form of Parkinsonism called X-linked dystonia Parkinsonism (XDP)^64^. The mechanism by which altered function of such a broadly important TF — TAF1 is part of the transcription preinitiation complex — can result in this particular phenotype has remained mysterious; our analysis, by suggesting that TAF1 has a particularly strong regulatory role in substantia nigra, sheds light on this question.

Our tissue-specific results also suggest new master-regulatory relationships for further exploration (see Figure 4). For example, while we recovered the known roles of CEBPB in liver^65^ and blood^66^, we also detected a robust tissue-specific activating role for this TF in pancreas, where it was our top result. It has been pointed out that, though CEBPB is not a classic pancreatic TF^65^, it is expressed in pancreatic beta cells specifically when they are under metabolic stress^65^; our result therefore suggests an *in vivo* pancreatic regulatory role for this TF that may be more easily detected using our eQTL-based analysis than using model systems that do not necessarily incorporate this environmental stimulus. Similarly, in addition to the roles we detected for HNF4A and HNF4G in liver, we also detected robust tissue-specific activating effects for both TFs in stomach, a less well-known association that has only recently been suggested^67,68^. We also identified a robust tissue-specific activating role for MAFF in skeletal muscle. This is interesting because, while the regulatory role of this TF in muscle is not well-studied, its expression is increased by an order of magnitude in muscle tissue after exercise^69^. MAFF is typically considered a transcriptional repressor, and indeed we observed a negative association between MAFF and activating histone marks in our previous analysis of molecular QTL in blood; the positive association we observe here therefore suggests that MAFF’s function in skeletal muscle may differ from its function in other tissues, perhaps via tissue-specific recruitment of an as-yet uncharacterized transcriptional activator. Finally, we also identified a robust tissue-specific negative role for CTCF in putamen (a brain tissue) and a robust tissue-specific activating role for the same TF in tibial artery. While CTCF is known to be capable of both repressive activity via insulation^47,48^ and activating activity^70^, this analysis suggests that its repressive/activating role varies meaningfully from tissue to tissue.

In addition to demonstrating how signed LD profile regression can be used to dissect transcriptional regulation in individual tissues, our results also demonstrate how our method can offer insights into aspects of transcriptional regulation that are not tissue-specific. For example, the transcription factor YY1 is a pioneer factor that has recently attracted considerable interest^71–74^. This TF has been theorized via detailed experimental work to mediate enhancer-promoter interaction^75^, but of the thousands of genes differentially expressed following *YY1* knockdown, approximately as many increase as decrease in their expression level^75^, presumably due to downstream regulatory cascades. In contrast, our analysis, which due to its use of eQTLs is able to focus primarily on cis-regulatory effects rather than downstream responses, shows a robust, predominantly activating role for YY1 across 25 tissues.

### Analysis of 46 diseases and complex traits

We applied signed LD profile regression to 46 diseases and complex traits with an average sample size of 289,617, including 16 traits with publicly available summary statistics and 30 UK Biobank traits for which we have previously publicly released summary statistics computed using BOLT-LMM v2.3^76^ (see URLs and Table S8). We first tested each of our 382 TF binding annotations for a directional effect on each of the 46 traits in turn (Table 1a and Table S9). For each significant association that we detected, we then evaluated 10,325 gene sets from the Molecular Signatures Database^23,24^ (MSigDB; see URLs) for enrichment among the genomic regions driving the association (controlling for LD and co-localizing genes; see Online Methods), in order to better understand the transcriptional programs mediating the association (Table 1b and Table S10).

**Table 1.**
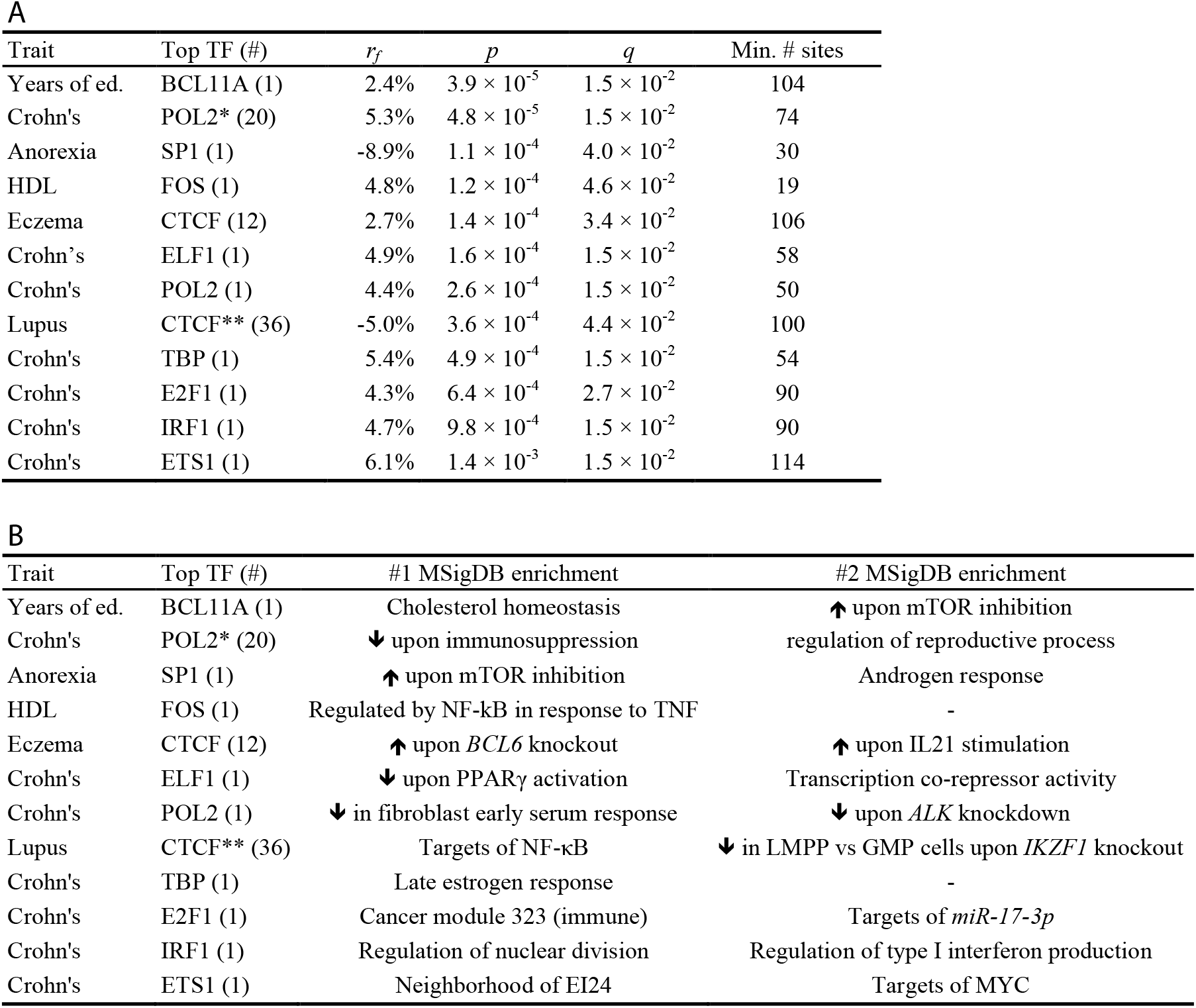
Independent TF-trait associations from analysis of diseases and complex traits using signed LD profile regression. For each of 12 independent associations at per-trait FDR<5% after pruning correlated annotations (*R*^2^ ≥ 0.25), we report the associated trait; the TF of the most significant annotation and the number of correlated annotations with significant associations; (a) the estimated functional correlation *r_f_*, P-value, q-value, and minimum number of TF binding sites contributing to the association; (b) the top two significant MSigDB gene-set enrichments among loci driving the association. Linked TFs producing significant associations: (*) TAF1, TBP; (**) RAD21. See Table S10 for full gene set names and enrichment q-values (all <5 × 10^−2^). LMPP: lymphoid-primed pluripotent progenitor; GMP: granulocyte-monocyte precursor.

Our analysis yielded 77 significant annotation-trait associations at a per-trait FDR of 5%, spanning six diseases and complex traits (see Figure 5 and Table S9a). (Following standard practice, we report per-trait FDR, but we estimated the global FDR of this procedure to be 9.4%, which is larger than the per-trait FDR of 5%; see Online Methods). The 77 significant associations represent 12 independent TF-trait associations after pruning correlated annotations (Table 1; see Online Methods). Of the 12 independent TF-trait associations, 9 involve an auto-immune disease as the phenotype, representing a 4.3x enrichment (*p* = 1. 9×10^−5^ using one-sided binomial test) and providing additional evidence for the relevance of TF binding to these phenotypes in particular^77^. To verify empirically that our results were not driven by confounding due to directional effects of minor alleles, we re-analyzed our data using an alternate set of 382 annotations defined using the same set of SNPs with non-zero effects but with the directionality of effect determined by minor allele coding rather than predicted TF binding, for SNPs in the bottom quintile of the MAF spectrum. This analysis yielded only 4 significant annotation-trait associations at per-trait FDR < 5%, implying that minor-allele-driven confounding is unlikely to explain our results. (Due to the small number of associations relative to the number of traits, these 4 minor-allele associations have a global FDR of 92.9% after accounting for 46 traits.) Furthermore, none of these 4 minor-allele associations overlapped with our set of 77 significant associations (see Online Methods and Table S9b). We also examined, for each annotation, the estimated covariance between the GWAS summary statistics and the signed LD profile in each of 300 independent genomic blocks, finding agreement with the genome-wide direction of association in 59% of the blocks on average across our 12 independent associations, and in 85% of the blocks with estimated covariances of large magnitude (see Figure S8). We used a related approach to compute a lower bound on the number of independent TF binding sites contributing to each association (Table 1a; see Online Methods). This lower bound ranged from 19 to 114, with an average value of 74 across the 12 independent TF-trait associations.

**Figure 5.**
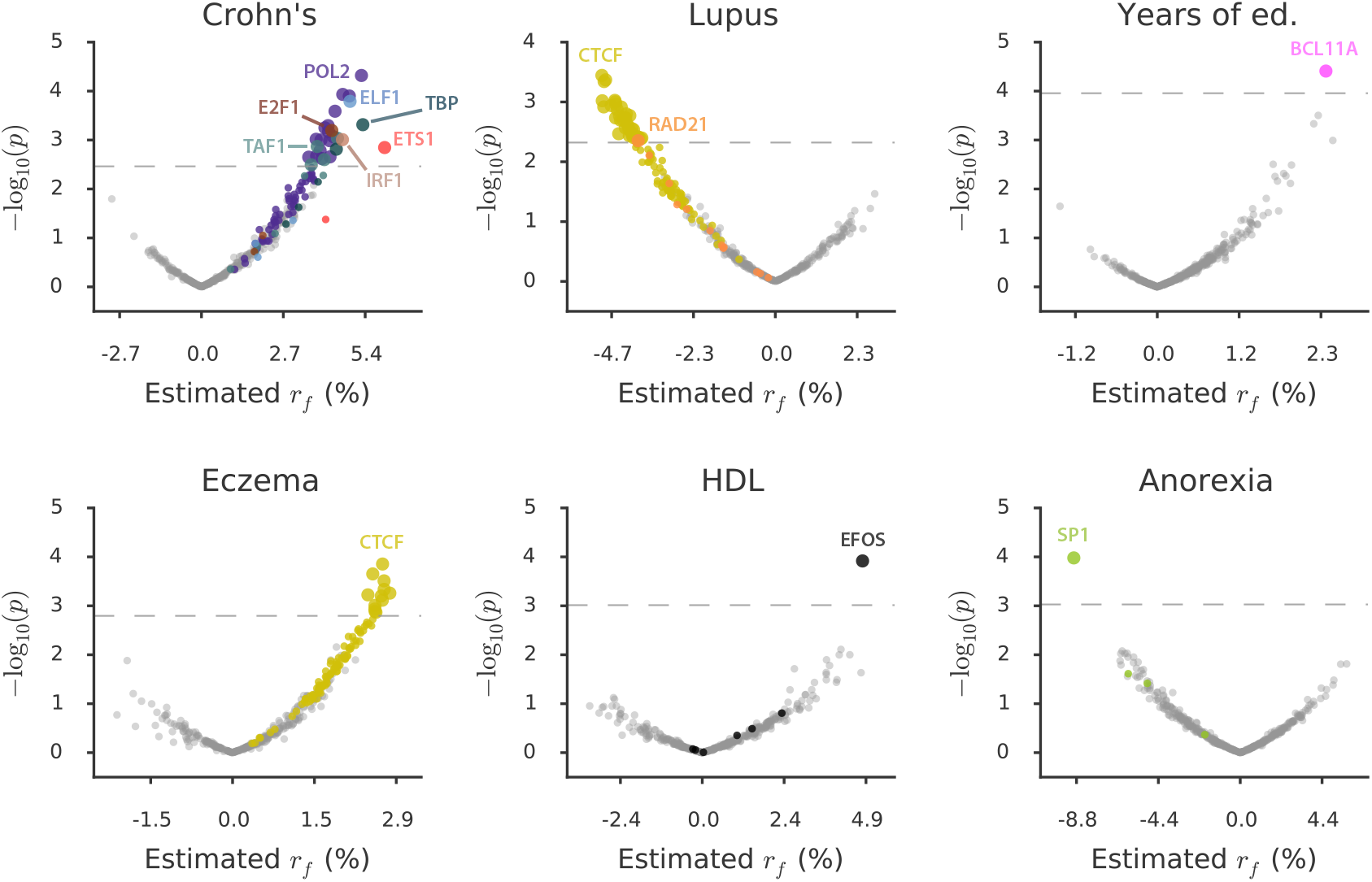
Analysis of diseases and complex traits using signed LD profile regression. For each disease or complex trait with at least one significant result, we plot –log_10_ (*p*) against estimated effect size for each of the 382 annotations analyzed. Points are colored by TF identity, with TFs with no significant associations for the trait colored in gray. Larger points denote significant results. The number of significant results for each trait is: Crohn’s, 26; Lupus, 36; Years of education, 1; Eczema, 12; HDL, 1; Anorexia, 1. Numerical results are reported in Table S9a.

Some of the TF-trait associations that we detected deepen our understanding of well-established associations or support and refine emerging theories of disease (Figure 6 and Table S11), while others were previously unknown (Figure 7 and Table S12). We begin by discussing three selected TF-trait associations that build on previous knowledge (Figure 6). First, we detected a positive association between genome-wide binding of BCL11A and years of education (see Figure 6a) that aligns well with existing evidence from educational attainment GWAS^78^, rare-variant studies of intellectual disability^79–82^, and experimental work showing that heterozygous knockout of *Bcllla* in mice leads to microcephaly and cognitive impairment^82^. (Additionally, our fine-mapping of the BCL11A GWAS locus using CAVIAR^83^ identified a putatively causal SNP in an intron of the *BCL11A* gene; see Table S13.) This association thus represents a case in which our method provides insight into the mechanism of a known relationship: specifically, we establish that BCL11A causes intellectual disability via binding *throughout the genome* — likely modulating (in cis) genes comprising a transcriptional program relevant to brain function or development — rather than regulation of a single key disease gene (see Discussion). Furthermore, our MSigDB gene-set enrichment analysis of the genomic regions driving the genome-wide signal allows us to characterize this putative transcriptional program. Specifically, we observed that these genomic regions are significantly enriched for an mTOR signaling gene set as well as for genes involved in cholesterol metabolism (see Figure 6a and Table S10). Regarding the mTOR gene set, the *MTOR* gene is itself an intellectual disability gene that has been intimately linked to brain development^84,85^. Regarding the cholesterol metabolism gene set, the brain contains approximately 25% of the body’s cholesterol (mostly as a component of the myelin sheaths that surround axons)^86,87^ with defects in brain cholesterol metabolism being linked to central nervous system disease^88,89^, and BCL11A has recently been shown to influence (and be influenced by) lipid levels^90–92^. Furthermore, the cholesterol metabolism and mTOR gene-set enrichments may be related, as mTOR has been linked to cholesterol metabolism^93^, including in the developing brain^94^. Because these gene-set enrichments characterize the genes putatively regulated in cis by BCL11A to affect brain function, this raises the possibility that mTOR exerts part of its effect on intellectual disability either by regulating or acting in concert with BCL11A to influence cholesterol metabolism in the developing brain.

**Figure 6.**
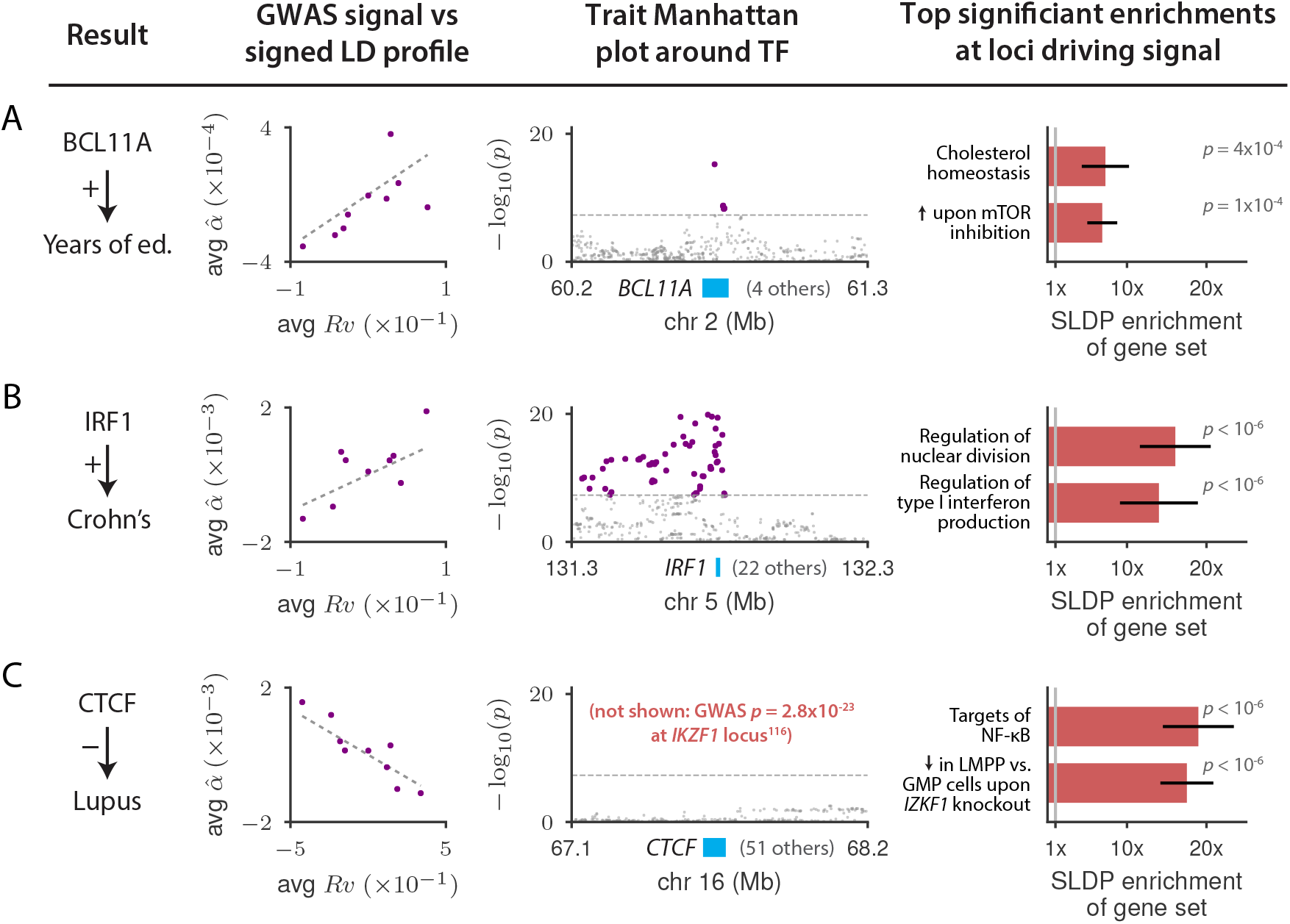
Highlighted TF binding-complex trait associations that either provide biological insight into established genetic associations or refine emerging theories. For each of (a) BCL11A-Years of education, (b) IRF1-Crohn’s disease, (c) CTCF-Lupus associations, we display plots of the marginal correlation 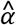 of SNP to trait versus the signed LD profile *Rν* of the annotation in question, with SNPs collapsed into bins of 4,000 SNPs and a larger bin around *Rν* = 0; Manhattan plots of the trait GWAS signal near the associated TF; and the top two significant MSigDB gene-set enrichments among the loci driving the association, with error bars indicating standard errors. Numerical results are reported in Table S11. LMPP: lymphoid-primed pluripotent progenitor; GMP: granulocyte-monocyte precursor.

**Figure 7.**
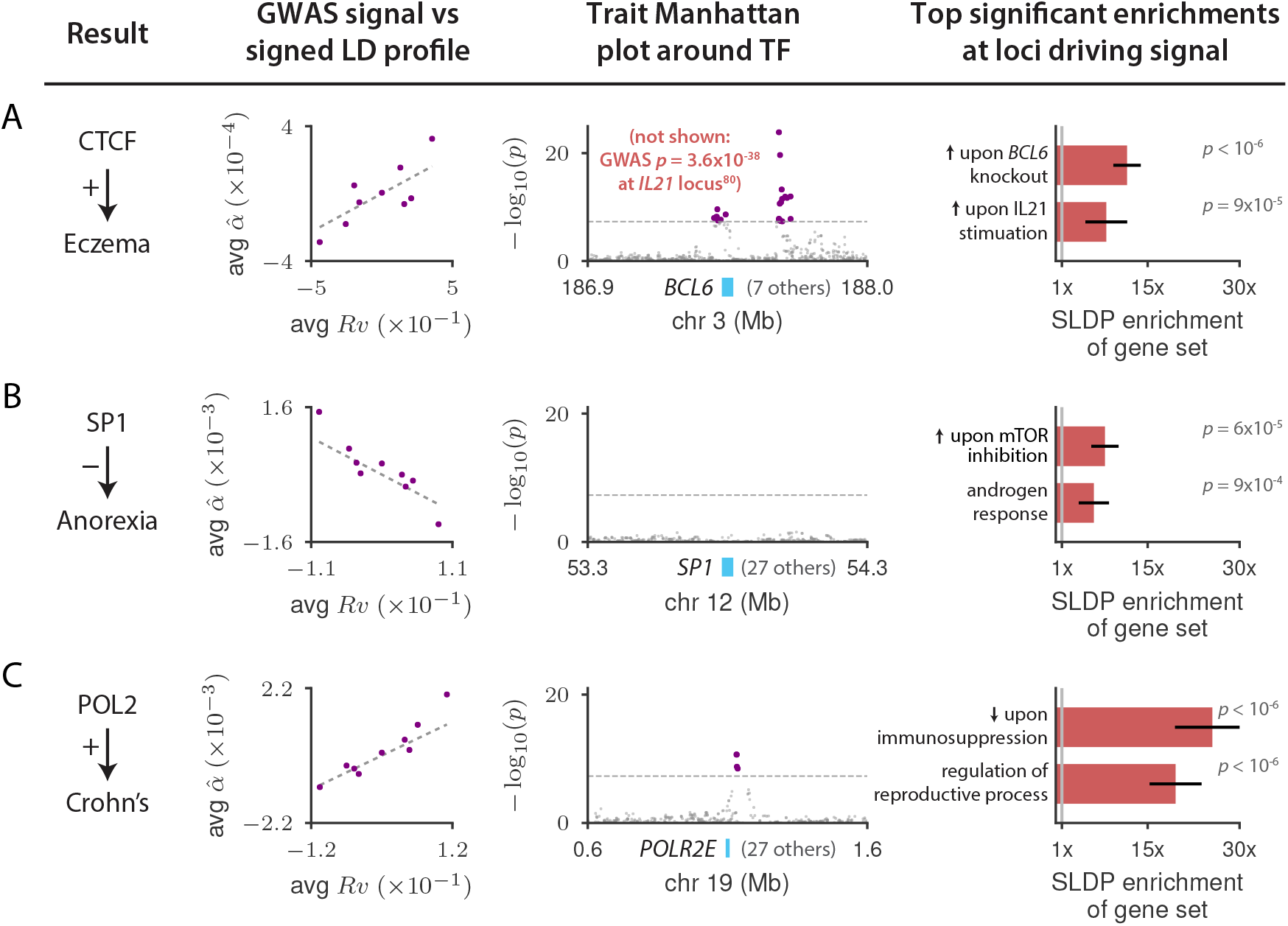
Highlighted previously unknown TF binding-complex trait associations. For each of (a) CTCF-Eczema, (b) SP1-Anorexia, (c) POL2-Crohn’s disease, we display plots of the marginal correlation *a* of SNP to trait versus the signed LD profile *Rν* of the annotation in question, with SNPs collapsed into bins of 4,000 SNPs and a larger bin around *Rν* = 0; Manhattan plots of the trait GWAS signal near the associated TF or, in the case of CTCF-Eczema, the *BCL6* gene (see main text; there is no GWAS peak at *CTCF*); and the top two significant MSigDB gene-set enrichments among the loci driving the association, with error bars indicating standard errors. Numerical results are reported in Table S12.

Second, we detected a positive association between genome-wide binding of interferon regulatory factor 1 (IRF1) and Crohn’s disease (CD) (see Figure 6b), a case in which existing GWAS evidence has been suggestive but not conclusive. Although *ÍRF1* is located inside a locus associated with CD and inflammatory bowel disease in multiple GWAS^95–97^ (one of the earliest CD associations^98^), this locus remains mysterious (it is known as the *“IBD5* locus”^99^, named after the *IBD5* gene). Strong LD makes it challenging to determine which variant(s) are causal, and high gene density at the locus (23 protein-coding genes within 500kb of *IRF1*) complicates the task of determining which gene is affected by any putative causal variant, resulting in several genes^95,100^ (including *IBD5^101^*) being previously nominated as potentially causal. For example, a recent large-scale fine-mapping study^102^ narrowed down the causal signal to a set of 8 SNPs including rs2188962, an eQTL for *SLC22A5* in immune and gut epithelial cells^22,102^ but also for *IRF1* in blood^35^. The transcriptome-wide association study (TWAS) approach^103^ for prioritizing genes has also been inconclusive: it assigns highly significant scores to both *ÍRF1* and *SLC22A5*, as well as five other genes at the locus whose predicted expression is positively associated to CD.^104,105^ Our result provides genome-wide evidence for a genuine causal link between IRF1 and CD that, unlike single-locus approaches, is not fundamentally limited by LD and pleiotropy near the *ÍRF1* gene (see Discussion). The top results in our MSigDB gene-set enrichment analysis strengthen our finding: the regions driving this association are most significantly enriched for genes involved in production of type I interferon and for genes involved in regulation of nuclear division (see Figure 6b and Table S10), matching well-known regulatory roles of IRF1^106,107^ and suggesting that IRF1 may affect CD via production of type-I interferon and concomitant cell-cycle regulation. We note that several other TF-trait associations from our analysis implicate a causal gene at an established GWAS locus, including ELF1-CD and ETS1-CD, with gene-set enrichments suggesting connections to existing CD drugs and to the role of autophagy in CD pathogenesis, respectively (see Table 1 and Supplementary Note).

Third, we detected a negative association between genome-wide binding of CCCTC-binding factor (CTCF) and risk of systemic lupus erythematosus (see Figure 6c) that supports an emerging theory of disease. Although there exists anecdotal evidence linking CTCF binding to lupus risk at a few isolated loci^108–110^, these results are once again susceptible to the effects of LD and pleiotropy, whereas our approach is able to provide stronger evidence for a causal relationship using genome-wide evidence involving TF binding at many concordant loci (at least 100; see Table 1a). We note that we do not observe a GWAS signal for lupus at the *CTCF* locus. This could be because the *CTCF* gene is under strong selective constraint (probability of loss-of-function intolerance^111^ = 1.00, greater than *99.9%* of genes), and/or because of the small sample size of the lupus GWAS. This association therefore demonstrates that signed LD profile regression can yield gene-disease associations in cases when GWAS is under-powered near the gene in question due to selection or small sample size. Our MSigDB gene-set enrichments shed additional light on this relationship: though CTCF has many diverse regulatory functions throughout the genome, the genomic regions driving the CTCF-Lupus association are most significantly enriched in immune gene sets, with the two strongest enrichments being targets of NF-ĸB and genes differentially expressed between two different stages of myeloid differentiation under knockout of the gene *IKZF1* (but not in the presence of IKZF1) (see Figure 6c and Table S10). The latter gene-set enrichment, because it pertains to genes putatively regulated in cis by CTCF, suggests a detailed mechanism whereby IKZF1 (itself a transcription factor) regulates or acts in concert with CTCF to activate a broader transcriptional program that opposes myeloid differentiation and reduces lupus risk. This hypothesis makes three predictions, each of which has evidence in the literature and/or publicly available data that we analyzed: (i) It predicts that IKZF1 affects Lupus risk; indeed, the *IKZF1* gene lies inside a Lupus GWAS locus^112,113^. (ii) It predicts that CTCF affects myeloid development; indeed, CTCF has been experimentally shown to slow myeloid differentiation^114,115^. (iii) It predicts that IKZF1 modulates CTCF activity; indeed, we determined that IKZF1 has ChIP-seq peaks in the vicinity of the *CTCF* promoter^116,117^ (see Table S14), consistent with a direct effect of IKZF1 binding on *CTCF* expression, and IKZF1 ChIP-seq peaks have also been shown to be enriched for the CTCF motif^118^, suggesting that these two TFs may also work in concert at binding sites throughout the genome. Thus, the association between CTCF binding and lupus that we detected, together with the associated MSigDB gene-set enrichments, enhances our understanding of the lupus GWAS signal at the *IKZF1* locus by providing evidence for *IKZF1* as the causal gene (out of 7 total protein coding genes within 500kb); suggests a mechanism to explain the effect of *IKZF1* on lupus; and proposes a regulatory relationship between *IKZF1* and *CTCF* that unifies disparate molecular evidence for the effects of both of these genes on myeloid development and ties them jointly to lupus risk.

We next discuss three selected TF-trait associations that were previously unknown (Figure 7). First, we detected a positive association between genome-wide binding of CTCF and eczema (see Figure 7a) that contrasts with the negative association that we detected between CTCF and lupus. The association with eczema exhibits gene-set enrichments that are very different from lupus. Moreover, the top two significant MSigDB gene-set enrichments for CTCF-Eczema are convergent: genes up-regulated in Treg cells upon knockout of the inflammatory regulator *BCL6;* and genes up-regulated in response to stimulation by the immune signaling molecule IL21, which is a known regulator of BCL6 activity^119,120^ (see Figure 7a and Table S10). As with the CTCF-Lupus example, these enrichments suggest a detailed cascade that we hypothesize to modulate eczema risk: IL21 signaling regulates BCL6, which in turn regulates or acts in concert with CTCF to activate a broad transcriptional program that increases eczema risk. This hypothesis makes three predictions: (i) It predicts that BCL6 modulates CTCF activity; indeed, we determined that BCL6 has many binding sites near the *CTCF* promoter in publicly available ChIP-seq data^121–124^ (see Table S15). (ii),(iii) It predicts that IL21 and BCL6 each affect eczema risk; indeed, the *IL21* and *BCL6* genes each fall in eczema GWAS loci^76,125,126^ (in each case along with 7 other protein-coding genes within 500kb). Thus, the association between CTCF binding and eczema that we detected nominates causal genes at two different existing eczema GWAS loci and provides a parsimonious mechanism for how both causal genes exert their effect on eczema via a regulatory cascade that drives a CTCF-mediated transcriptional program.

Second, we detected a negative association between genome-wide binding of SP1 and anorexia (Figure 7b), a heritable trait for which no single locus reaches genome-wide significance in the GWAS data that we analyzed^127^. SP1 levels have been shown observationally to correlate negatively with psychiatric conditions such as bipolar disorder^128^ and schizophrenia^129,130^ (which is significantly positively genetically correlated with anorexia^131^), but this association has not been shown to be causal and has not previously been observed in GWAS of psychiatric traits. Our MSigDB gene-set enrichment results for this association yielded significant enrichments for an androgen response gene set and an mTOR signaling gene set (see Figure 7a and Table S10). (Years of education, for which an mTOR signaling gene-set was also among the top two MSigDB enrichments, is also significantly positively genetically correlated with anorexia^131^; the median rank of the top-scoring mTOR gene set across the 10 other independent TF-complex trait associations was 1,123, of 10,325 MSigDB gene sets tested.) The androgen response result is intriguing given the sex-imbalanced nature of this phenotype^132^. The mTOR signaling result is noteworthy given the well-established connections between mTOR, caloric restriction, and growth^133^; it also raises the possibility that a link between SP1 and mTOR could explain prior observations that SP1 can be regulated by insulin levels^134,135^, modulate expression in the hypothalamus of the appetite regulator POMC^136,137^, and play a role in the induction of leptin following insulin-stimulated glucose metabolism in adipocytes^138^. In addition, mTOR has been shown to play an important role in androgen signaling^139^, suggesting a potential unification of these two signals.

Third, we detected a positive association between binding of RNA polymerase II (POL2) and Crohn’s disease (CD) (Figure 7c). This association is surprising given the very broad role of POL2 throughout the genome. However, our MSigDB gene-set enrichments shed some light on the biology underlying this association, with many significant enrichments in immune and immune-related gene sets (see Table S10). In particular, the top two significant gene sets are genes down-regulated upon immunosuppression and genes involved in cell-cycle regulation (see Figure 7c and Table S10). Because of the central role of POL2 in gene transcription, these results suggest that there may exist a large set of immune- or proliferation-related genes whose increased expression contributes to CD risk. Indeed, CD is an auto-immune disease, and it has been hypothesized that increased expression is a prominent component of many immune responses since it can be enacted more quickly than decreased expression^140–142^. Furthermore, acute inflammation has been associated in observational studies with CD onset^143,144^, and recent experimental work^145^ has shown that the acute inflammatory response in mice is greatly attenuated by non-specific inhibition of the general-purpose transcriptional machinery containing POL2. Our result potentially links these two findings, providing evidence that the observational association between acute inflammation and CD is causal and suggesting that there exists a polygenic liability for acute inflammation that acts via increased transcription of a large set of immune- or proliferation-related genes and contributes to CD risk. To better understand the POL2-CD association, we investigated whether any of the 14 genes comprising the RNA polymerase II protein complex lie inside a CD GWAS locus. We identified a CD GWAS peak located 28kb from one of these genes, *POLR2E.* This locus is quite gene-dense (28 protein-coding genes within 500kb; 3 protein-coding genes within 28kb), and a recent large-scale CD fine-mapping effort^102^ was unable to nominate any gene as potentially causal. Thus, our POL2-CD association also nominates a potential causal gene for the CD GWAS association at this gene-dense locus.

We provide additional discussion of other TF-trait associations in the Supplementary Note.

## Discussion

We have introduced a method, signed LD profile regression, for identifying genome-wide directional effects of signed functional annotations on diseases and complex traits. We first applied this method, in conjunction with 382 annotations describing predicted effects of SNPs on TF binding, to 12 molecular traits in blood (average *N* = 149) and confirmed that it recovers classical aspects of transcriptional regulation, including the pro-transcriptional effect of RNA polymerase and activating TFs as well as associations between chromatin modifiers and their respective chromatin marks. We then applied the method to gene expression eQTLs in 48 GTEx tissues from the GTEx consortium (average *N* = 214), yielding 2,330 significant annotation-tissue expression associations representing 651 distinct TF-tissue expression pairs, 30 of which showed strong evidence of tissue specificity. These included many previously unknown associations that support emerging theories of disease in less well-studied tissues and new tissue-specific master-regulatory relationships. Finally, we applied the method to 46 diseases and complex traits (average *N* = 289,617), identifying 77 annotation-trait associations, representing 12 independent TF-trait associations. Some of these findings confirm previously well-established associations, others provide insight into known GWAS loci containing the associated TF (in addition to other protein-coding genes), and others have not been detected in prior GWAS. Because the detected associations involve genome-wide TF binding, they implicate broad disease-relevant transcriptional programs. Our characterization of these programs via gene-set enrichment analyses using gene sets from MSigDB^23,24^ yielded detailed hypotheses about disease mechanisms that in several cases mechanistically link existing GWAS loci and disparate molecular evidence into a parsimonious mechanism mediated by the associated TF.

Our method differs from unsigned GWAS enrichment methods^1–7^ by assessing whether there is a systematic genome-wide correlation between a signed functional annotation and the (signed) true causal effects of SNPs on disease, rather than assessing whether a set of SNPs have large effects on a disease without regard to the directions of those effects. A substantial advantage of this approach is reduced susceptibility to confounding: for example, an unsigned GWAS enrichment for binding of an immune TF could indicate a causal role for that TF in the associated disease, or could instead be a side effect of a generic enrichment among cell-type-specific regulatory elements in immune cells^5^. Unsigned enrichments can also be complicated by LD, as functional elements in LD with binding sites of a TF may contribute to its enrichment if not properly modeled^5^. In contrast, if alleles that increase binding of the TF tend to increase disease risk and alleles that decrease binding of the TF tend to decrease disease risk, the set of potential confounders is smaller because a confounding process has not only to co-localize in the genome with binding of the TF but also to have the property that alleles that increase the process have a consistent directional effect on binding of the TF.

Our method differs from existing single-locus GWAS methods^11,12,103^ in that it enables stronger statements about causality and mechanism. Regarding causality, this is because a consistent genome-wide directional effect of SNPs predicted to affect TF binding due to sequence change (across a large set of TF binding sites; see Table 1a) is less susceptible to pleiotropy, LD, and allelic heterogeneity^103,105^. The robustness of our method to these potential confounders is also greater than that of genetic correlation and Mendelian randomization^131^ (MR) analyses, which can be confounded by reverse causality and pleiotropic effects^146–148^ (and which would scale poorly because they would require TF ChIP-seq in many individuals for every TF/cell-type pair studied). The reason that our method is not confounded by reverse causality is that each of our annotations is produced in a cell population that is isogenic and therefore does not have variance in genetic liability for any trait. In other words, our annotations provide ideal instrumental variables for the effect of TF binding on the trait of interest because they are created not by naively correlating SNPs with TF binding but rather by examining the effect of each SNP on local DNA sequence.

Regarding mechanism, our method sheds light on the question of whether TFs affect traits via coordinated regulation of gene expression throughout the genome^149^ (a “genome-wide” model) or via regulation of one or a small number of key disease genes^150^ (a “local” model). Since the associations we find involve a consistent net direction of effect of TF binding on a trait throughout the genome, they cannot be explained by a local model and therefore represent evidence for the existence of transcriptional programs and their relevance to complex traits.

This is of basic interest, but it also has therapeutic relevance: if a TF causally affects a trait but the TF is not druggable due to its nuclear localization or large DNA- and protein-binding domains^151,152^, then the local model suggests targeting a downstream gene, whereas the genome-wide model instead suggests targeting an upstream regulator since the causal link between TF and trait is mediated through a large number of downstream genes. (We emphasize that a significant result for our method does not imply that all binding events of the TF in question affect disease via activation of a single transcriptional program; rather, it implies that there exists a program that is widespread enough that we observe its effect on disease in a large number of locations in the genome; see Table 1a and Figure S8.) Moreover, as we have shown, the genome-wide nature of the putative transcriptional programs identified by our method allows us to characterize and interpret these programs by aligning them with existing gene sets, leading in some cases to detailed mechanistic hypotheses.

We note that although we constructed our predicted TF binding annotations using the neural-network predictor Basset^19^, there exist many other effective methods for making such signed predictions^1,13–16,18,153,154^ and many other data sets on which to train them^155–157^. In an initial effort to assess these, we repeated our analyses of molecular traits in blood, gene expression in 48 GTEx tissues, and 46 diseases and complex traits using annotations generated via three other approaches: 382 annotations generated using the DeepSEA neural-network predictor^15^ applied to the same ENCODE ChIP-seq data that we analyzed using Basset; 184 annotations generated using the Basset predictor trained on a larger but noisier set of meta-analyzed ChIP-seq data from the Gene Transcription Regulation Database^4^ (GTRD) followed by our Basset QC procedures; and 276 annotations generated using position-weight matrices (PWMs) from the *Homo sapiens* Comprehensive Model Collection^156^ (HOCOMOCO), which are based in part on data from the GTRD (see Online Methods). Results are reported in Tables S16, S17, and S18, respectively, and summarized in Table S19. For the 382 DeepSEA annotations, we obtained results similar to our primary set of 382 Basset annotations, including replication of many of our top results (see Figures S9 and S10 and Table S16); intriguingly, we also determined that the concordance between signed LD profile regression results using Basset and DeepSEA was greater than the concordance between Basset and DeepSEA at the level of annotations (see Figure S11), suggesting that the signal that is shared between the predictions made by the two methods is indeed biological. The DeepSEA annotations produced fewer significant associations in total (see Table S19), although this comparison was restricted to annotations passing our Basset QC procedures, including a filter on Basset prediction accuracy (see Figure S10). The 184 GTRD annotations produced fewer significant annotations than either set of annotations created using ENCODE data, though they did identify new associations, especially in GTEx eQTL data (see Tables S17 and S19). For the 276 PWM-based annotations from HOCOMOCO, we again observed correlation between results using PWMs and results using Basset (see Figures S12 and S13), though this correlation was weaker than the correlation between the DeepSEA results and the Basset results. We identified fewer significant associations overall using the PWM-based annotations than we did using the more sophisticated neural-network based annotations (see Tables S18 and S19), providing evidence that the latter methods can provide a scientifically meaningful increase in performance.

Our method could be used to link disease to biological processes beyond TF binding. For example, sequence-based models can also produce signed predictions of DNase I hypersensitivity^14,15,19^, histone modifications^15,19^, splicing^16,158^, and transcription initiation^159^. Additionally, allele-specific molecular assays, massively parallel assays, and CRISPR screens are increasingly yielding high-resolution experimental information about the effects of genetic variation on gene expression^29,45,160–163^ as well as cellular processes such as growth^164–166^ and inflammation^167^. Finally, perturbational differential expression experiments can yield signed predictions for the relationships of genes to a variety of biological processes such as drug response^168^, immune stimuli^169^, and many others^170^. Though converting such data to signed functional annotations will require care, doing so could allow us to leverage them to make detailed statements about disease mechanism.

We note several limitations of signed LD profile regression. First, though our results are less susceptible to confounding due to their signed nature, they are not immune to it: in particular, our method cannot distinguish between two TFs that are close binding partners and thus share sequence motifs, and it likewise cannot distinguish between binding of the same TF in different cell types, as the resulting annotations could be highly correlated. Second, although we have shown our method to be robust in a wide range of scenarios, we cannot rule out the possibility of un-modeled directional effects of minor alleles on both trait and TF binding as a confounder; however, our empirical analysis of real traits with minor-allele-based signed annotations suggests that directional effects of minor alleles are very unlikely to explain our results (see Table S9b). Third, our results are limited by the quality of the annotations we are able to produce. For example, TF binding is easier to measure in open chromatin and so it may be the case that our annotations for activating TFs are more representative of underlying biology — and therefore better powered — than our annotations for repressing TFs. Fourth, our method is not well-powered to detect instances in which a TF affects trait in different directions via multiple heterogeneous programs. Fifth, the effect sizes of the associations to diseases and complex traits that we report are small in terms of the estimated values of *r_f_*, which range in magnitude from 2.4% to 8.9% (recall that *r_f_* is analogous to a genetic correlation; see Table S9a), although signals of this size for predicted TF binding could be indicative of much stronger associations, e.g., with true TF binding, TF expression, TF phosphorylation, or TF binding in specific subsets of the genome. We further note that the magnitude of the signals that we detect is commensurate with the very small number of SNPs in our annotations. Specifically, 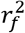 divided by the proportion of SNPs in an annotation quantifies how much heritability the signed TF binding signal that we detect explains as compared to the total heritability explained by a random set of SNPs of the same size. This ratio is as large as 3.5x (see Table S9c), implying that our signed TF binding signals can account — in a signed fashion — for substantial trait heritability relative to the proportion of SNPs. Sixth, we used annotations constructed using ChIP-seq data from cell lines, which is non-ideal both because chromatin dynamics in cell lines do not necessarily match those in real tissue and because cell lines often have structural duplications and deletions that complicate sequence-based analysis of TF binding. We note, however, that though these difficulties reduce our power and so are promising topics for future work, they would not be expected to introduce false positives into our results due to the signed nature of our analysis. Seventh, our annotations are constructed by testing each minor allele in the context of the reference genome and separately from variation at all other SNPs, rather than taking into account potential non-linear interactions between nearby SNPs^1^; this too is a source of reduced power but not increased false positive rates. Eighth, the interpretability of our MSigDB gene-set enrichment analysis is limited by the potential for distinct gene sets to have overlapping membership as well as the possibility for co-expressed genes to be in the same gene sets more often than expected by chance; however, we believe this is somewhat ameliorated by that fact that we treat blocks of genes together in our empirical null (see Online Methods). Ninth, though we detected many significant associations overall, there were many diseases and complex traits, including schizophrenia, height, and blood cell traits, for which we did not detect any significant associations using our TF annotations. We believe that three factors may contribute to this: (i) As we observed here and as others have noted as well^77^, auto-immune traits appear to have a stronger association to TFs than other traits, at least for the TFs on which we have systematic, high-quality ChIP-seq data, and these traits comprised only 8 out of 47 (17%) of the diseases and complex traits in our study; it may be that genome-wide directional effects of these TFs are not as prominent a mechanism for other traits. (ii) We construct our annotations by annotating all SNPs in the ChIP-seq peaks for the TF in question; it could be that in many cases these annotations represent multiple opposing or unrelated transcriptional programs, and that restricting them to more specific sets of SNPs would reveal additional genome-wide directional effects. (iii) Genome-wide directional effects may be contingent on annotations constructed using data generated in the “correct” cellular context (beyond the narrow set of cell lines analyzed in this paper). It is possible that additional signed TF-trait associations will be identified as higher-quality functional data sets become more available and molecular hypotheses become more detailed.

Despite these limitations, signed LD profile regression is a powerful new way to leverage functional genomics data to draw causal and mechanistic conclusions from GWAS about both diseases and underlying cellular processes.

## Acknowledgements

We thank C de Boer, L Dicker, J Engreitz, T Finucane, N Friedman, R Gumpert, M Kanai, S Kim, X Liu, M Mitzenmacher, J Perry, S Reilly, D Reshef, S Raychaudhuri, A Schoech, P Sabeti, R Tewhey, O Troyanskaya, P Turley, O Weissbrod, J Zhou, and the CGTA discussion group for helpful discussions. This research was conducted using the UK Biobank Resource under Application #16549 and was supported by US National Institutes of Health grants U01 HG009379, R01 MH101244 and R01 MH107649. L.P. is supported by National Institutes of Health award R00HG008399. R.P.A. is supported by NSF IIS-1421780. Computational analyses were performed on the Orchestra High Performance Compute Cluster at Harvard Medical School, which is partially supported by grant NCRR 1S10RR028832-01.

## URLs

Signed LD profile regression: open-source software is available at http://www.github.com/yakirr/sldp

Plink2: https://www.cog-genomics.org/plink2/

BLUEPRINT consortium data: ftp://ftp.ebi.ac.uk/pub/databases/blueprint/blueprint_Epivar/qtl_as/QTL_RESULTS/

TWAS weights for NTR data: https://data.broadinstitute.org/alkesgroup/FUSION/WGT/NTR.BLOOD.RNAARR.tar.bz2 GTEx eQTL data: https://www.gtexportal.org/home/datasets MSigDB data: http://software.broadinstitute.org/gsea/msigdb GTRD data: http://gtrd.biouml.org/

HOCOMOCO motif data: http://hocomoco11.autosome.ru/

## Online Methods

### Signed LD profile regression

We first describe the method intuitively, then present a formal derivation and discuss other technical details.

### Intuition

Our method for quantifying directional effects of signed functional annotations on disease risk, signed LD profile regression, relies on the following intuition. Suppose there are *M* SNPs and we are given a signed functional annotation, specified by a length-*M* vector *ν*, with a directional linear effect on disease risk. For example, *ν* might be a vector whose *m*-th entry is the effect of SNP *m* on binding of some TF. If we knew the length-*M* vector *β* of the true causal effects of the same SNPs on a trait, we could simply regress *β* on *ν* to evaluate whether there is a non-trivial signed association across SNPs m between *ν_m_* and *ν_m_*. In reality, we cannot do this because we do not observe *β*; instead we observe a vector, denoted 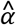, of GWAS summary statistics describing the marginal correlation of every SNP to our trait of interest. This vector differs from *β* because it includes both causal and tagging effects, plus statistical noise. Specifically, it can be shown mathematically that, in expectation, 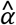 will equal the matrix-vector product *Rβ* where *R* is the *M×M* LD matrix. Therefore, just as *β* would be proportional to *ν* in the presence of a signed effect, 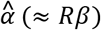 would likewise be proportional to *Rν*, which is a vector capturing each SNP’s aggregate tagging of the signed annotation. This means that instead of regressing *β* on *ν* (which is impossible since we do not observe *β*), we can regress 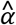 on *Rv*. We call the vector *Rv* the *signed LD profile* of *ν*, and thus our method is called signed LD profile regression. The remainder of our technical material is oriented toward i) weighting this regression to achieve optimal power, ii) being able to efficiently perform the required computations, iii) determining the proper way to test the null hypothesis of no signed effect, and iv) controlling for potential confounding due to directional effects of minor alleles.

### Model and estimands

Let *M* be the number of SNPs in the genome. We assume a linear model:

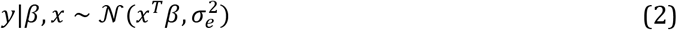

where 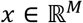 and 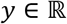 are the standardized genotype vector and phenotype, respectively, of a randomly chosen individual from some population, 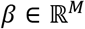 is a vector of true causal effects of each SNP on phenotype, and 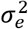 represents environmental noise. Given a signed functional annotation 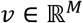, we then model

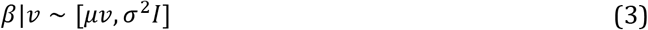

where the scalar *μ* represents the genome-wide directional effect of *ν* on *β*, *σ*^2^ represents other sources of heritability unrelated to *ν*, and the notation [·,·] is used to specify the mean and covariance of the distribution without specifying any higher moments.

Though we can estimate *μ*, its value depends on the units of the annotation and the heritability of the trait. Because of this, we focus instead on the *functional correlation r_f_*, which re-scales *μ* to be dimensionless and is defined as

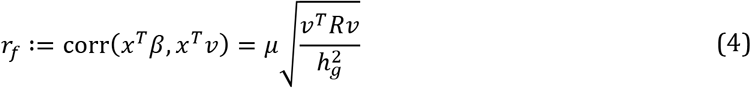

where 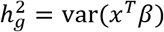 is the SNP-heritability of the phenotype and 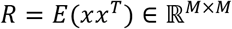 is the (signed) population LD matrix of the genotypes. (Note that *r_f_* can also be defined as a correlation between *β* and *ν*; this definition is approximately equivalent in expectation under our random effects model, provided *ν^τ^Rν* ≈ |*ν*|^2^.) We additionally estimate 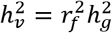, the total phenotypic variance explained by the signed contribution of *ν* to *β*, as well as 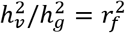. For annotations with small support, these quantities are expected to be small in magnitude. To see this, notice that 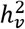 cannot exceed the total (unsigned) phenotypic variance explained by SNPs with nonzero values of *ν*. It follows that 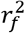 cannot exceed the proportion of (unsigned) SNP-heritability explained by SNPs with non-zero values of *ν*. For more detail on the model and estimands, see the Supplementary Note.

### Main derivation

Let *X* ∈ ℝ^*N×M*^ be the genotype matrix in a GWAS of *N* individuals, with standardized columns, and let *Y* ∈ ℝ^*N*^ be the phenotype vector. In the Supplementary Note, we show that under the above model the following identity approximately holds:

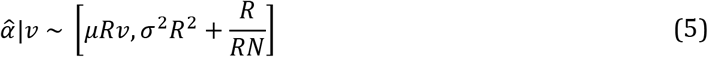

where 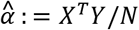 is a vector whose *m*-th entry contains the marginal correlation of SNP *m* to the phenotype and *R* ∈ ℝ^*M×M*^ is the population LD matrix. Equation (1) from the main text can be derived from Equation (5) by re-scaling v so that *ν^τ^Rν* = 1, then substituting for *μ*.

We call *Rν* the *signed LD profile* of *ν*. Equation (5) means that we can estimate *μ* by regressing 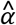 on the signed LD profile using generalized least-squares with *Ω*: = *σ*^2^*R*^2^ + *R/N* as the inverse weight matrix. It can be shown that if a) all causal SNPs are typed, b) sample size is infinite, and c) *R* is invertible, this method is equivalent to estimating *β* via 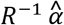 and then regressing this estimate on *ν* to obtain *μ*, which is the optimal regression-based approach in that setting. Note that because we generate P-values for hypothesis testing empirically (see below), we are guaranteed that our generalized least-squares scheme will remain well-calibrated even if our estimate of the matrix *Ω* is inaccurate due to, e.g., mis-match between the reference panel and the study population. Once we have estimated *μ*, we re-scale this estimate to yield an estimate of *r_f_* and other estimands of interest. For more detail on derivations and computational considerations, see the Supplementary Note.

### Null hypothesis testing

To test the null hypothesis *H*_0_: *μ* = 0 (or, equivalently, *H*_0_: *r_f_* = 0), we split the genome into approximately 300 blocks of approximately the same size with the block boundaries constrained to fall on estimated recombination hotspots^171^. We then define the null distribution of our statistic as the distribution arising from independently multiplying *ν* by one independent random sign per block. We perform this empirical sign-flipping many times to obtain an approximation of the null distribution and corresponding P-values. Our use of sign-flipping ensures that any true positives found by our method are the result of genuine first-moment effects; if in contrast we estimated standard errors using least-squares theory or a re-sampling method such as the jackknife or bootstrap, our method might inappropriately reject the null hypothesis only because the variance of *β* is higher in parts of the genome where *Rν* is large in magnitude. This would make our method susceptible to confounding due to unsigned enrichments, as might arise from the co-localization of TF binding sites with enriched regulatory elements such as enhancer regions. Additionally, the fact that we flip the signs of SNPs in each block together ensures that our null distribution preserves any potential association of our annotation to the LD structure of the genome. In choosing how many blocks to use for this procedure, we took into account that i) the fewer blocks we use the fewer assumptions we make about LD structure and the faster we can compute P-values, and ii) the more blocks we use the higher the precision of the P-values that we can obtain. Our choice to use 300 blocks is a compromise between these two considerations.

### Controlling for covariates and the signed background model

Given a signed covariate *u* ∈ ℝ^*M*^, we can perform inference on the signed effect of *ν* conditional on *u* by first regressing *Ru* out of 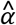 and out of *Rν* using the generalized least-squares method outlined above, and then proceeding as usual with the residuals of 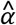 and *Rν*. This can be done simultaneously for multiple covariates *u*.

Unless stated otherwise, all analyses in this paper are done controlling in this fashion for a “signed background model” consisting of 5 annotations *u*^1^,…, *u*^5^, defined by

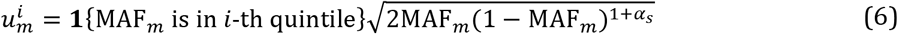

where MAF_*m*_ is the minor allele frequency of SNP *m* and *α_s_* is a parameter describing the MAF-dependence of the signed effect of minor alleles on phenotype. Based on the literature on MAF-dependence of the unsigned effects var(*β_m_*), we set *a_s_* = –0.3^172^.

### 382 TF annotations

We downloaded every ChIP-seq and DNase I hypersensitivity experiment in ENCODE and trained the sequence-based predictor of peak presence/absence, Basset^19^, to jointly predict each downloaded track on a set of held-out genomic segments. (We included tracks other than TF binding tracks because training predictions using all tracks slightly improved prediction accuracy for the TF binding tracks.) After training the joint predictor, we retained the predictions for every TF binding track for which a) the number of SNPs in the set of ChIP-seq peaks with non-zero difference in Basset predictions between the major and minor allele was at least 5,000 in our 1000G reference panel, and b) Basset’s estimated area under the precision-recall curve (AUPRC) was at least 0.3. This yielded a set of 382 TF ChIP-seq experiments. For each experiment, we constructed an annotation via

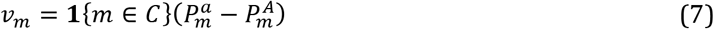

where *C* is the set of SNPs in the ChIP-seq peaks arising from the experiment, 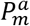 is the Basset prediction for the 1,000 base-pair sequence around SNP m when the minor allele is placed at SNP *m*, and 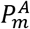 is the Basset prediction for the 1,000 base-pair sequence around SNP *m* when the major allele is placed at SNP *m*. (We always used the minor allele as the reference allele in both our TF binding annotations and our GWAS summary statistics.)

### Simulations

All simulations were carried out using real genotypes from the GERA cohort^27^ (*N* = 47,360). The set of *M* = 2.7 million causal SNPs was defined as the set of very well imputed SNPs (INFO ≥ 0. 97) that had very low missingness (< 0.5%) and non-negligible MAF (MAF ≥ 0.1%) in the GERA data set, and were represented in our 1000G Phase 3 European reference panel^146,173^.

### Null simulations

For the simulations in Figure 1a, we simulated 1,000 independent null phenotypes with the architecture 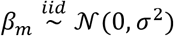 with 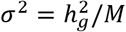 and 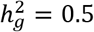. For each phenotype, we computed GWAS summary statistics using plink2^174^ (see URLs), adjusting for 3 principal components as well as GERA chip type as covariates. For each of our 382 TF annotations, we then ran signed LD profile regression on each of these 1,000 phenotypes, yielding a set of 382,000 P-values. For the simulations in Figure 1b, we simulated 1,000 independent traits in which each trait had an unsigned enrichment for a randomly chosen annotation: after choosing an annotation *ν*, we set 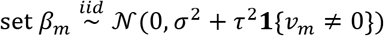 where *σ*^2^ and *τ*^2^ were set to achieve 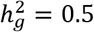 and a 20x unsigned enrichment for the SNPs with non-zero values of *ν*. We then computed summary statistics as above and ran signed LD profile regression to assess v for a genome-wide directional effect. This procedure yielded 1,000 P-values. For the simulations in Figure 1c, we simulated 1,000 independent phenotypes with a directional effect of minor alleles: we set 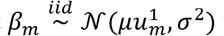 where 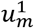 is non-zero if SNP m is in the bottom quintile of the MAF spectrum of the GERA sample and 0 otherwise, as in the signed background model. We set *μ* such that 10% of heritability would be explained by this directional effect, and then set *σ*^2^ to achieve 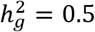. We then computed summary statistics as above and ran signed LD profile regression to assess for a directional effect of each of our 382 annotations on each of the 1,000 phenotypes, yielding a set of 382,000 P-values. Finally, we repeated the same computation but running signed LD profile regression without the 5-MAF-bin signed background model to obtain an additional set of 382,000 P-values.

### Causal simulations

For the simulations in Figure 2, we fixed a representative annotation v (binding of IRF4 in GM12878), and simulated traits using 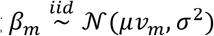, with *μ* set to achieve *r_f_* = {0,0.005,0.01,…,0.05} and *σ*^2^ set to achieve 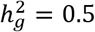 in each case. For each value of ry, we simulated 100 independent traits, computed summary statistics using plink2, and then ran each of the methods under consideration using the annotation *ν*.

### Analysis of molecular traits in blood

We downloaded BLUEPRINT consortium QTL data for gene expression, H3K4me1, H3K27ac, and methylation in three different blood cell types with sample sizes of *N* = 158, 165, and 125 for monocytes, neutrophils, and T cells, respectively^21^ (see Table S4 and URLs). For each of the 3 gene expression traits, we constructed one summary statistics vector 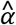 by setting

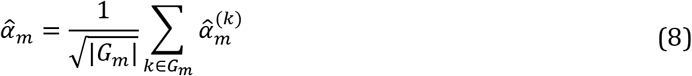

where *G_m_* is the set of all genes within 500kb of SNP *m*, and 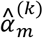 is the marginal correlation of SNP *m* to the expression of gene *k*. Assuming independence of expression across genes this is analogous to a fixed-effects meta-analysis across genes at every SNP to determine that SNP’s effect on aggregate expression, though our results do not rely on this theoretical characterization because of the empirical, signed nature of our null hypothesis testing procedure. Since in practice gene expression is not independent across genes, the scale of the resulting vector 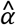 is arbitrary. Therefore, we placed all such vectors on the same scale by scaling them so that they have an estimated SNP-heritability of 0.5. (This scaling step only affects the regression weights used by signed LD profile regression.) Applying the same procedure to the two histone marks and to methylation in addition to gene expression yielded a total of 12 sets of summary statistics (see Table S4). We ran signed LD profile regression using each of our 382 TF annotations for each of these 12 traits. We obtained results at FDR < 5% using the Benjamini-Hochberg procedure^175^ within each of the 12 traits (see discussion of Benjamini-Hochberg versus other alternatives below), and reported the union of significant results across cell types for each trait. We determined the top 100 associations to display in Figure 3a by choosing the significant associations with the highest estimated values of *r_f_*.

For our replication analysis, we used expression array-based whole blood eQTL data from the NTR^35^, which we obtained by downloading the set of TWAS weights^103^ computed for that data set (see Table S4 and URLs). We then proceeded as above. We note, however, that because TWAS weights were only available for genes with a significantly heritable cis-expression in NTR, we only had data for 2,454 genes compared with 15,023 – 17,081 genes for the BLUEPRINT traits, thereby lowering our power in this analysis.

### Enrichment analysis for activating TFs

For each TF represented in our annotations, we queried the UniProt database^31^ to establish whether the TF was (unambiguously) “activating”, “ambiguous”, or (unambiguously) “repressing” (see Results). To estimate whether the set of significant positive signed LD profile associations with gene expression were enriched for (unambiguously) “activating” TFs compared to the set of annotations as a whole, we conducted a one-sided binomial test. To account for the correlated nature of our annotations, we assumed independence only among distinct TFs but not among distinct annotations for the same TF. We used the same scheme to test for enrichment of (unambiguously) “activating” TFs among the positive associations detected by signed LD profile regression in our analysis of histone marks.

### Analysis of gene expression across 48 GTEx tissues

We downloaded GTEx v7 eQTLs for all 48 tissues for which data were available and processed them using the same procedure described for the blood molecular traits, resulting in one vector of summary statistics per GTEx tissue (see Table S6 and URLs). We ran signed LD profile regression using each of our 382 TF annotations for each of these tissues. We obtained results at FDR< 5% using the Benjamini-Hochberg procedure^175^ within each of the 48 tissues (see discussion of Benjamini-Hochberg versus other alternatives below).

### Conditional analysis for tissue-specific effects

We obtained a set of eQTL summary statistics for a fixed-effect meta-analysis across the GTEx tissues from ref.^176^ and processed these via the procedure described above into a single vector 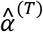. For each tissue *t*, we then residualized 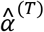 out of the vector 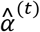 of eQTL data for tissue *t* to obtain a residualized vector 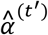. This simply amounts to subtracting a scalar multiple of 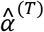 from 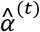, with the scalar determined to remove as much signal as possible from 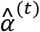. For each significant association between an annotation *a* and a vector 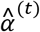 from our main GTEx analysis, we then compared the p-value of that association to the p-value obtained for the association between *a* and the residualized vector 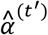, declaring as tissue-specific any association for which the latter was at least as significant as the former. For cases in which a P-value for association to either 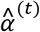 or 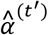 was ≤ 10^−5^ (one order of magnitude greater than the maximal resolution of of our empirical null hypothesis testing procedure), we replaced that p-value by a closed-form p-value computed by constructing a z-score out of the estimated value of *r_f_* and its jackknife-based standard error.

### Assessment for concordance with absolute expression levels in GTEx tissues

We obtained raw gene expression levels across the GTEx samples as in ref.^177^ and filtered both the raw expression levels and our 382 TF binding annotations to the set of 68 TFs that were represented in both data sets. (This procedure excluded, e.g., POL2, which does not correspond to a single gene.) For each of the 34 GTEx tissues *t* in which we detected significant association(s) among these 68 TFs, we then computed *p_t_*, the proportion of the significant TFs in that tissue with a median transcripts per million (TPM) value greater than 5 across the GTEx samples for that tissue (following ref.^75^), and *q_t_*, the proportion of the remaining TFs in that tissue with a median TPM value greater than 5 across the GTEx samples for that tissue. Figure S7 contains a plot of *p_t_* against *q_t_* across tissues *t*. To evaluate the significance of the trend across tissues that *p_t_* > *q_t_*, we compared 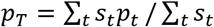 to 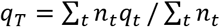 where *s_t_* and *n_t_* are the numbers of TFs with significant associations and without significant associations, respectively, in tissue *t*. We then rejected the null hypothesis that *p_T_ ≤ q_T_* using a one-sided two-sample z-test for difference in means.

### Analysis of 46 diseases and complex traits

We applied signed LD profile regression to 46 diseases and complex traits with an average sample size of 289,617, including 16 traits with publicly available summary statistics and 30 UK Biobank traits for which we have publicly released summary statistics computed using BOLT-LMM^76^ (see Table S8 and URLs). We ran signed LD profile regression using each of our 382 TF annotations for each of these traits. We obtained results at per-trait FDR < 5% using the Benjamini-Hochberg procedure^175^. We chose to use the Benjamini-Hochberg procedure rather than more sophisticated procedures such as the Storey-Tibshirani procedure^178^ because the latter procedure, while more powerful, is more difficult to analyze in a multi-trait setting (see below) and controls FDR more noisily when applied in situations with only hundreds (rather than thousands) of tests.

### MSigDB gene-set enrichment analysis of results on diseases and complex traits

We downloaded all 10,325 MSigDB gene sets, which are organized into eight distinct tranches based on their origin, from the MSigDB online portal. We also downloaded a set of LD blocks in Europeans derived from estimated recombination hotspots^171^ and converted each gene set into a length-1693 vector s with one entry per LD block whose *i*-th entry equaled the number of genes from the set that are present in the *i*-th LD block. We then converted each significant signed LD profile regression association between an annotation *ν* and a trait summary statistics vector 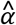 into a length-1693 vector *q* whose *i*-th entry equaled the covariance between 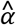 and the signed LD profile *Rν* within the *i*-th LD block. To assess the signed LD profile result for enrichment of a gene-set vector s, we computed a weighted mean of the *q_i_* whose weights were given by *s*. That is, we computed 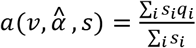 The idea is that if the LD blocks in which *s* is large correspond to the LD blocks in which the signed LD profile regression signal is the strongest, the weighted mean *a* should be large in magnitude and have the same sign as the overall signed LD profile regression association. We assess this via an empirical null distribution constructed by permuting the LD blocks to obtain “shuffled” versions of *s* and *q*. This enrichment method is more conservative than ordinary gene-set enrichment methods for two reasons. First, by permuting only LD blocks and not genes, it accounts for correlations induced by LD as well as co-regulation of nearby genes and gene overlap in the genome. Second, because a significant signed LD profile regression association cannot arise as a result of a strong signal in only one genomic location, this method is more robust to outliers and cannot, e.g., produce a rejection simply because of a very strong signal at just one gene. In comparison to gene-set enrichment methods for GWAS data, this method also has the advantage that it will not cause gene sets containing large genes to produce signals of enrichment. Separately from null hypothesis testing, we computed heuristic standard errors for use in Figures 6 and 7 by computing the closed-form standard deviation of 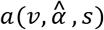 assuming that the *s_i_* are fixed and the *q_i_* are i.i.d.

To quantify effect size, we computed a fold-enrichment by dividing 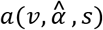 by the average value of *q* at LD blocks containing no genes. That is the enrichment is defined as 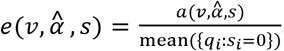. This quantity *e* is the number reported in Figures 6 and 7.

We conducted our hypothesis test for gene-set enrichment for each of our 77 significant TF-complex trait associations against each of the 10,325 MSigDB gene sets. For every TF-complex trait association and every tranche of gene-sets from MSigDB, we assessed significance at FDR< 5% using the Benjamini-Hochberg procedure^175^. This detected 6,379 significant enrichments in total (0.8% of all 795,025 tests conducted). We ranked these enrichments by q-value, except for the 15 enrichments whose p-values were less than the resolution of our empirical null hypothesis testing procedure, which we ranked by fold-enrichment.

### Estimation of global FDR for complex trait analysis

When many traits are analyzed, per-trait FDR control does not imply global FDR control. This is because in the case of a completely null trait, the guarantee of FDR control does not imply that there will never be any rejections but rather only that there will be a non-zero number of rejections at most 5% of the time. Therefore, if enough null traits are analyzed the set of results may be contaminated by these spurious findings. In the case of independent tests (i.e., uncorrelated annotations) with FDR controlled by the Benjamini-Hochberg procedure, this can be taken into account^179^ and the global FDR can be approximated using the formula

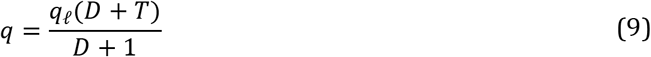

where *q* is the estimated global FDR, *q_ℓ_* is the per-trait FDR, *D* is the observed total number of discoveries at per-trait FDR *q_ℓ_*, and *T* is the number of traits. This correction is based on the intuition that for a null trait with independent tests, the Benjamini-Hochberg procedure behaves very similarly to a Bonferroni correction, and so the expected number of rejections per null trait is approximately *q_ℓ_*, and the expected number of rejections for *T* null traits would be approximately *q_ℓ_T*.

Applying this correction to our results yields a global FDR estimate of 7.9%. However, since our annotations are dependent, this estimate can be anti-conservative. To see this, imagine a null trait with 100 perfectly correlated tests. The Benjamini-Hochberg procedure will give more than zero rejections only 5% of the time, but whenever it rejects it will yield 100 rejections rather than 1. Therefore, the expected number of rejections is not 0.05 but rather 5. We heuristically corrected for this using the intuition that under dependent tests, the expected number of false discoveries in a null stratum is not *q_ℓ_* but rather *q_ℓ_* times the number of tests conducted per single “independent” test. We estimated the number of independent tests as in the GWAS literature, by simulating 1,000 independent null traits with a heritability of 0.5, testing each trait against our 382 annotations, and asking for what S we see at least one p-value ≤ 0.05/*S* in approximately 5% of the 1,000 null traits. This procedure gave us *S* = 250. We then estimated the global FDR using the equation

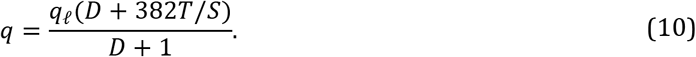

This yielded the reported global FDR of 9.4%.

### Pruning 77 significant associations to 12 independent signals

To prune our set of 77 significant associations to a set of approximately independent results, we used the following iterative greedy approach for each trait: we chose the pair of associations whose annotations had the most strongly correlated signed LD profiles, removed the annotation with the less significant p-value, and repeated until no annotations in the result set had signed LD profiles that were correlated at *R*^2^ > 0.25. We used correlation between signed LD profiles rather than between the annotations themselves because, since our method regresses the summary statistics on the signed LD profile rather than the raw annotation, correlation between signed LD profiles most accurately represents the correlation between the test statistics for the two annotations. Grouping the results by TF identity gives similar results (13 distinct TF-trait associations as opposed to 12 independent TF-trait associations; see Table S20).

### Analysis of diseases and complex traits with annotations corresponding to directional effects of minor alleles

We constructed an alternate set of 382 annotations as follows. For each of the 382 ChIP-seq experiments represented by a set of peaks *C*, we set

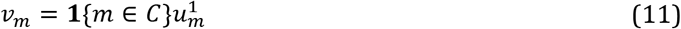

where *u*^1^ is the signed background annotation corresponding to SNPs in the bottom quintile of the MAF spectrum. We then used signed LD profile regression to test for association between each of these 382 annotations and each of our 46 traits, assessing significance as above.

### Estimation of lower bound on number of independent TF binding sites contributing to each association

We converted each of the 12 independent TF-trait associations reported in Table 1 into a vector *q* of length ~300 whose *i*-th entry equaled the covariance between the GWAS in question and the signed LD profile in question within the *i*-th of the ~300 independent genomic blocks used for our null hypothesis testing. For every threshold 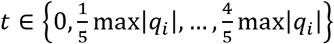, we then computed the number *K_t_* of the entries of *q* with magnitude at least *t*, as well as the number *S_t_* of those entries whose sign agreed with that of the genome-wide trend. Our estimated lower bound on the number of independent TF binding sites contributing to the association was then given by

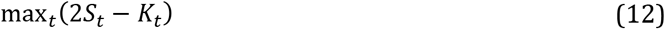

The intuition for this is that the distribution of the signs of the entries of *q* can be modeled as a mixture of a uniform distribution (for genomic chunks with no signal) and a distribution with all of its mass on the sign of the genome-wide trend (for genomic chunks with signal). The number of entries drawn from the latter distribution is what we seek to estimate. This is because it gives the number of independent genomic blocks contributing to the association, which is a lower bound on the number of independent TF binding sites contributing to the association (since a genomic block spanning 1/300-th of the genome may contain multiple independent TF binding sites). Estimating this number naively without thresholding yields the expression 2*S*_0_ – *K*_0_. However, this is an under-estimate in the presence of noise in *q*. We therefore repeat this argument considering only the subset of entries of *q* with magnitude at least t for a small number of thresholds t and retain the largest estimate.

### Creation of additional annotations using DeepSEA, GTRD, and HOCOMOCO

#### Creation of additional annotations using DeepSEA

For each of the 382 ENCODE TF ChIP-seq tracks used to generate our post-QC Basset annotations, we obtained predictions for the same track using the DeepSEA method from the authors of that method. We then created 382 new annotations using the same procedure used to generate the 382 Basset annotations (see Equation (7)). We analyzed each of these annotations against the blood molecular QTL, the GTEx eQTL, and the 46 diseases and complex traits; for results, see Table S16 and Figures S9 and S11. We also obtained the reported AUPRCs of Basset and DeepSEA on all 691 of ENCODE TF ChIP-seq tracks; these are compared in Figure S10.

#### Creation of additional annotations using GTRD

We downloaded all 482 of the meta-cluster tracks from the GTRD (see URLs) and trained Basset to predict these tracks jointly with the ENCODE tracks used to train our main Basset predictor. We created 482 annotations from these tracks using the same procedure used to generate the 382 (ENCODE) Basset annotations (see Equation (7)). Only 149 (31%) of these annotations passed our standard QC filter (Basset prediction AUPRC > 0.3 and at least 5,000 SNPs with nonzero annotation values). We analyzed each of these 149 annotations against the blood molecular QTL, the GTEx eQTL, and the 46 diseases and complex traits; for results, see Table S17.

#### Creation of additional annotations using HOCOMOCO

We downloaded the 402 core human mononucleotide TF binding PWMs from the HOCOMOCO database (see URLs). We filtered these 402 PWMs to those for which the TF in question had a ChIP-seq track among the 382 post-QC ENCODE TF binding tracks used to produce our main set of annotations. For each of the resulting 58 PWMs, we then created one new annotation for every matching ENCODE TF binding track by using the PWM to score SNPs inside the ChIP-seq peaks in the matching track. This resulted in 276 annotations.

To create an annotation from a PWM and an ENCODE TF binding track, we first computed a score *t*(*x*) for every SNP allele *x* via 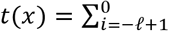 exp (pwm_*i*_(*x*)) where *ℓ* is the length of the PWM, and where pwm_*i*_ (*a*) is the PWM score given by the motif in question to the reference genome sequence with allele *x* substituted for the SNP in question and the first position of the PWM placed i bases before the SNP. (The PWM score of a sequence is the sum of the entries of the PWM specified by the bases comprising each position of the sequence^180^.) We then treated these scores as binding predictions and produced an annotation from them using the same procedure used to generate the 382 Basset annotations (see Equation (7)). We analyzed each of the resulting 276 annotations against the blood molecular QTL, the GTEx eQTL, and the 46 diseases and complex traits; for results, see Table S18 and Figures S12 and S13.

### Data availability

We have released all genome annotations we analyzed, as well as regression weight matrices for our 1000 genomes reference panel, at http://data.broadinstitute.org/alkesgroup/SLDP/.

### Code availability

Open-source software implementing our approach is available at http://www.github.com/yakirr/sldp. Code used to make all figures is available at http://www.github.com/yakirr/sldp-display.

